# Active shape programming drives *Drosophila* wing disc eversion

**DOI:** 10.1101/2023.12.23.573034

**Authors:** Jana F. Fuhrmann, Abhijeet Krishna, Joris Paijmans, Charlie Duclut, Suzanne Eaton, Marko Popović, Frank Jülicher, Carl D. Modes, Natalie A. Dye

**Author notes:** These authors contributed equally to this work (Jana F. Fuhrmann, Abhijeet Krishna).

## Abstract

How complex 3D tissue shape emerges during animal development remains an important open question in biology and biophysics. In this work, we study eversion of the *Drosophila* wing disc pouch, a 3D morphogenesis step when the epithelium transforms from a radially symmetric dome into a curved fold shape via an unknown mechanism. To explain this morphogenesis, we take inspiration from inanimate “shape-programmable” materials, which are capable of undergoing blueprinted 3D shape transformations arising from in-plane gradients of spontaneous strains. Here, we show that active, in-plane cellular behaviors can similarly create spontaneous strains that drive 3D tissue shape change and that the wing disc pouch is shaped in this way. We map cellular behaviors in the wing disc pouch by developing a method for quantifying spatial patterns of cell behaviors on arbitrary 3D tissue surfaces using cellular topology. We use a physical shape-programmability model to show that spontaneous strains arising from measured active cell behaviors create the tissue shape changes observed during eversion. We validate our findings using a knockdown of the mechanosensitive molecular motor MyoVI, which we find to reduce active cell rearrangements and disrupt wing pouch eversion. This work shows that shape programming is a mechanism for animal tissue morphogenesis and suggests that there exist intricate patterns in nature that could present novel designs for shape-programmable materials.

## Introduction

Epithelial tissues are sheets of tightly connected cells with apical-basal polarity that form the basic architecture of many animal organs. Deformations of animal epithelia in 3D can be mediated by external forces, either from neighboring tissue that induces buckling instabilities (e.g., [1-3]) or extracellular matrix that confines (e.g., [4]) or expands (e.g., [5]). Alternatively, local differences in mechanics at the apical and basal sides of the deforming epithelia itself can drive out-of-plane tissue shape changes (e.g., ventral furrow invagination in the *Drosophila* embryo (reviewed in [6]) and fold formation in *Drosophila* imaginal discs [7]).

Here, we describe a mechanism for generating complex 3D tissue shape involving tissue-scale patterning of in-plane deformations, analogous to the shape transformations of certain inanimate shape-programmable materials. These shape-programmable materials, like hydrogels and nematic elastomers, experience *spontaneous strains* where the local preferred lengths change in response to stimuli in a desired way [8, 9]. Globally patterned spontaneous strains can create a geometric incompatibility with the original shape, triggering specific, desired 3D deformations, such as the formation of a cone from a flat sheet [8, 10, 11]. Ideas from shape-programmability have already proved insightful to the understanding of differential growth-mediated plant morphogenesis [12, 13]. However, animal epithelia are more dynamic, changing cell shape and size, as well as rearranging tissue topology. As these behaviors cause in-plane changes in local tissue dimensions, the ingredients for shape-programmability are, in principle, present.

To test these concepts in animal morphogenesis, we quantify tissue shape changes and cell behaviors in the *Drosophila* wing disc during a 3D morphogenetic process called eversion (Fig. 1a). Through eversion, the wing disc proper, an epithelial mono-layer, undergoes a shape deformation in which the future dorsal and ventral surfaces of the wing blade appose to form a bi-layer and escape the overlying squamous epithelium called the peripodial membrane. After eversion, the wing disc begins resemble the final shape of the adult wing. This process is triggered by a peak in circulating levels of the hormone 20-hydroxyecdysone, analogous to an activator in shape programming. This complex tissue shape change is independent of forces external to the wing disc, as demonstrated by its ability to occur in explant culture [14]. The shape changes of the disc proper also cannot be fully explained by removal of the peripodial membrane or extracellular matrix and appear to be self-sufficient, involving active cellular processes [15-19].

**Fig. 1.**
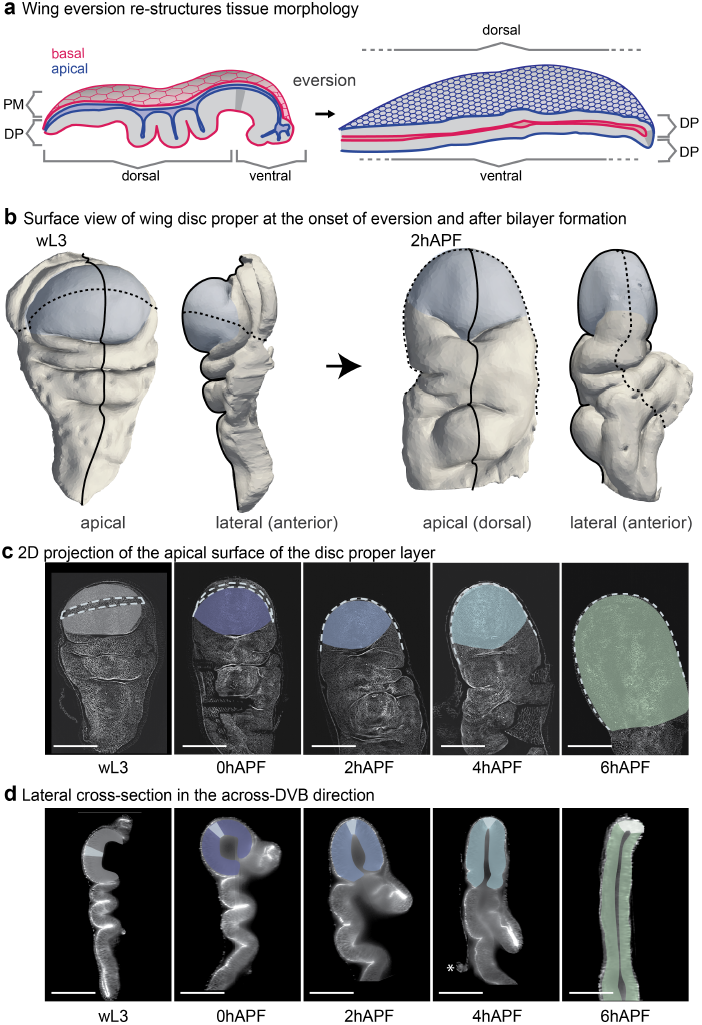
The wing pouch undergoes anisotropic curvature changes during eversion: **a**, Schematic cross-sections along the long axis of the wing disc before and after eversion. Before eversion, the wing disc resembles an epithelial sac with apical facing inwards. The tissue consists of the Disc Proper (DP), which is a folded, thick, pseudo-stratified mono-layer, and the Peripodial Membrane (PM), a thin squamous monolayer. After eversion, the PM is removed and the former pouch region of the DP forms the wing bilayer with apical facing out and dorsal and ventral on opposing sides. **b**, Example of a 3D segmentation of the DP in a head-on and side view before eversion (left, wL3) and after bilayer formation (right, 2hAPF). Pouch: blue; across-DVB: solid line; along-DVB: dashed line. **c**,**d**, Representative images for stages of eversion. Wing discs are labelled with Ecadherin-GFP. The pouch region is highlighted, colored by time. **c**, Projection view showing the dorsal side for early pupal stages and dorsal (down), ventral (up), and DVB for wL3. The position of the DVB is indicated with a dashed line. **d**, Across-DVB cross-section. The position of the DVB is indicated in white. Asterisk shows the rupture point of the PM, which gets removed around 4hAPF. Minimum 5 wing discs were analyzed for each time point; hAPF = hours after puparium formation; wL3 = wandering larval stage, 3rd instar; scale bars = 100 *µm*.

It has long been postulated that the eversion of wing (and leg) discs is achieved by in-plane cell behaviors that are organized by previously established cell morphology patterns [20-23]. Here, we test this hypothesis by systematic quantification and genetic perturbation of cell behaviors during eversion and demonstrate how cell behaviors contribute to tissue shaping using a physical model analogous to shape programming.

## 1 The wing pouch undergoes anisotropic curvature changes during eversion

We first sought to quantify the tissue shape changes happening during wing disc eversion. To this end, we explanted wing discs at fixed time intervals, from late larval stage (wL3) to 6 hours After Puparium Formation (hAPF). We imaged the wing discs using multi-angle light sheet microscopy and then reconstructed and analyzed the 3D image stack (Methods 7.3). In this way, we capture the complex 3D shape changes happening throughout the wing disc during eversion (Fig. 1b, Supplemental movie 1).

The most dramatic tissue shape changes can be seen in a central cross-section along the axis perpendicular to the dorsal ventral boundary (DVB), referred to as “across-DVB” (Fig. 1d, Extended Data Fig. S1d,f,g). We observe three main morphogenetic changes: the peripodial membrane is removed around 4hAPF, the deeply folded regions unfold, and the pouch undergoes a transition from a monolayer dome to a flat bilayer with a sharply folded interface. In the perpendicular plane, taken through DVB in the pouch (referred to as “along-DVB”), the tissue does not change as significantly, preserving curvature in this direction (Fig. 1c, Extended Data Fig. S1e-g).

We focus hereafter on the pouch region, as it undergoes the most complex shape change: starting as an almost radially symmetric dome and ending up in a curved-fold shape, with curvature increasing strongly in one axis (across-DVB) but not as much in the other (along-DVB) (Extended Data Fig. S1f,g). To test the hypothesis that in-plane cellular behaviors lead to 3D tissue shape change, we first build a shape-programmability model that can relate cellular behaviors to spontaneous strain. We then measure patterns of cellular behaviors in the wing pouch during eversion and use our model to test how they affect tissue shape change.

## 2 Programmable spring network as a model for epithelial morphogenesis

We developed a coarse-grained model of tissue shape changes, leveraging an analogy between tissue remodelling by internal processes and spontaneous strain-driven shape programming of nematic elastomers [24-26]. We use a double layer of interconnected programmable springs representing the apical surface geometry and the material properties of an epithelial sheet, including a bending rigidity introduced by the thickness of the double layer (Fig. 2a, Methods 7.9, 7.14). As an initial configuration, we use a stress-free spherical cap and then assign new rest lengths to the springs. In a continuum limit, this corresponds to introducing a spontaneous strain field 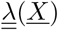 which depends on the spatial coordinates *<u>X</u>* Methods 7.10). To simplify notation, we write 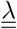 for 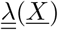 hereafter. To generate a final output shape, we quasi-statically relax the spring network (Methods 7.9, 7.10). As with conventional elastic strain tensors, 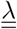 can be decomposed into isotropic (*λ*) and anisotropic 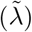 modes.

**Fig. 2.**
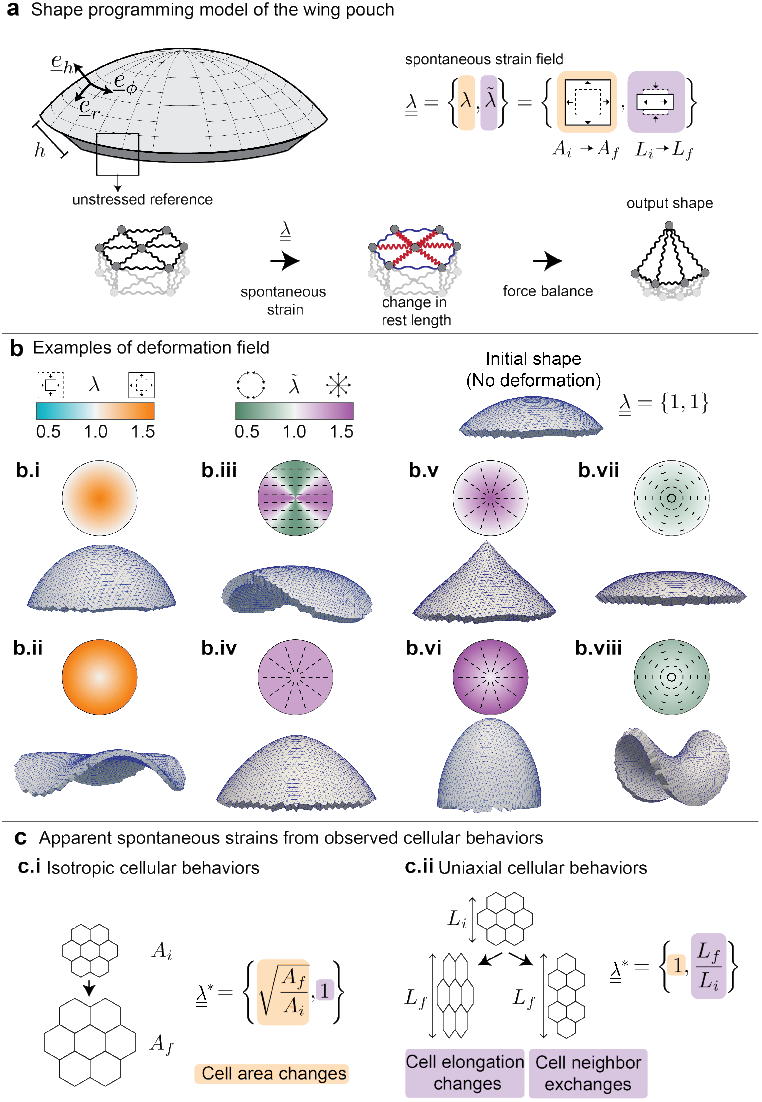
The programmable spring network relates cell behaviors to spontaneous strain to model epithelial morphogenesis: **a**, A thick spherical cap as a model for an epithelial tissue. We define a radial coordinate *r* and basis vectors <u>*e*</u>_*r*_, <u>*e*</u>_*ϕ*_, and <u>*e*</u>_*h*_. The thickness of the spring network h is constant everywhere and introduces a bending rigidity. The model tissue is an elastic medium implemented as a spring network with an initially stress-free state. We change the rest lengths of the springs by imposing a spontaneous strain field 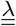 and allow subsequent relaxation to take a new 3D output shape. Top and bottom springs at any position in the lattice have their rest lengths updated by the same amount. The spontaneous strain field 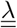 consists of an isotropic component λ and an anisotropic component 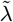. These components cause changes in area (*A*_i_ to *A*_f_) or area-preserving changes in shape (*L*_*i*_ to *L*_*f*_), respectively. **b**, Model realizations with simple patterns of spontaneous strains. For each realization, the input pattern of spontaneous strains is displayed above, with the magnitude of strain encoded by color. For anisotropic strain 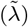, the bars indicate the orientation. Below is the output shape. In **b.i,ii**, we vary the isotropic contribution λ and keep 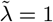, while in **b.iii-viii**, we vary 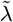 and keep λ = 1. We probe the model output from input linear radial gradients in λ or 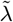, giving rise to cones with varying degrees of sharpness at the tip (**i,v,vi,vii**) or saddle shapes (**ii,viii**). Using a spatially homogeneous pattern of 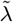, we observe an elongated spherical cap when patterned along a fixed direction (**iii**) and a blunt cone when patterned radially (**iv**). **c**, Schematics showing the calculation of apparent spontaneous strains from observed cellular behaviors. **c.i** For a patch of cells going from area of *A*_i_ to *A*_f_, we extract λ. **c.ii** A patch of cells undergoing anisotropic deformation due to cell elongation changes or neighbor exchanges causes the length scale in one direction to change from *L*_i_ to *L*_f_. From this change, 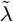 is extracted, while λ = 1, as there is no isotropic contribution.

We first wanted to understand how simple choices of spontaneous strain patterns induce a shape change in our model. A simple gradient of *λ*, for example, causes the spherical cap to balloon in the center or generate wrinkles at the periphery (Fig. 2b.i. and ii). Changing the directions and gradients of 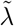 leads to elongation of the cap, increase in the curvature at the tip or even flattening of the curvature in the center, eventually leading to a saddle shape (Fig. 2b.iii - viii.).

We propose that cell behaviors can give rise to a spontaneous strain field, thereby shape programming the wing disc pouch and driving 3D shape changes during eversion. The strains measured from observed cell behaviors during eversion (referred to as observed strains, 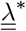) can be used to infer spontaneous strains. By using coarse-grained spontaneous strains, the topology of the spring network remains unchanged [27]. For the isotropic component of observed strain, we focus on cell area changes 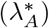, as cell division and cell death are minimal in the everting wing disc (Fig. 2ci) [14, 28, 29]. The anisotropic components of observed strain capture contributions stemming from both changes in cell elongation 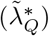 as well as from cell rearrangements 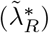 (Fig. 2cii).

Our model can therefore relate cell behaviors to spontaneous strains in order to understand resulting tissue deformations. We now investigate these quantities in the everting wing disc.

## 3 Topological tracking reveals spatial patterns of cell dynamics in the everting wing pouch

To examine cell behaviors, we first segmented apical cell junctions and plotted average cell area and cell elongation in space (Extended Data Fig. S2, Fig. 3a). From larval stages, we know that cell morphology and behaviors in the pouch are organized radially in the region outside of the dorsal ventral boundary (outDVB) and parallel to the boundary in the region closest to the dorsal ventral boundary (DVB) [30-33]. During eversion, we observe that cell shapes and sizes are patterned similarly. In early stages, cell area follows a radial gradient that disappears by the end of eversion (4hAPF) (Fig. 3a, Methods 7.7). Cell elongation exhibits a global nematic order through 4hAPF before disordering at 6hAPF (Fig. 3b, Methods 7.7).

**Fig. 3.**
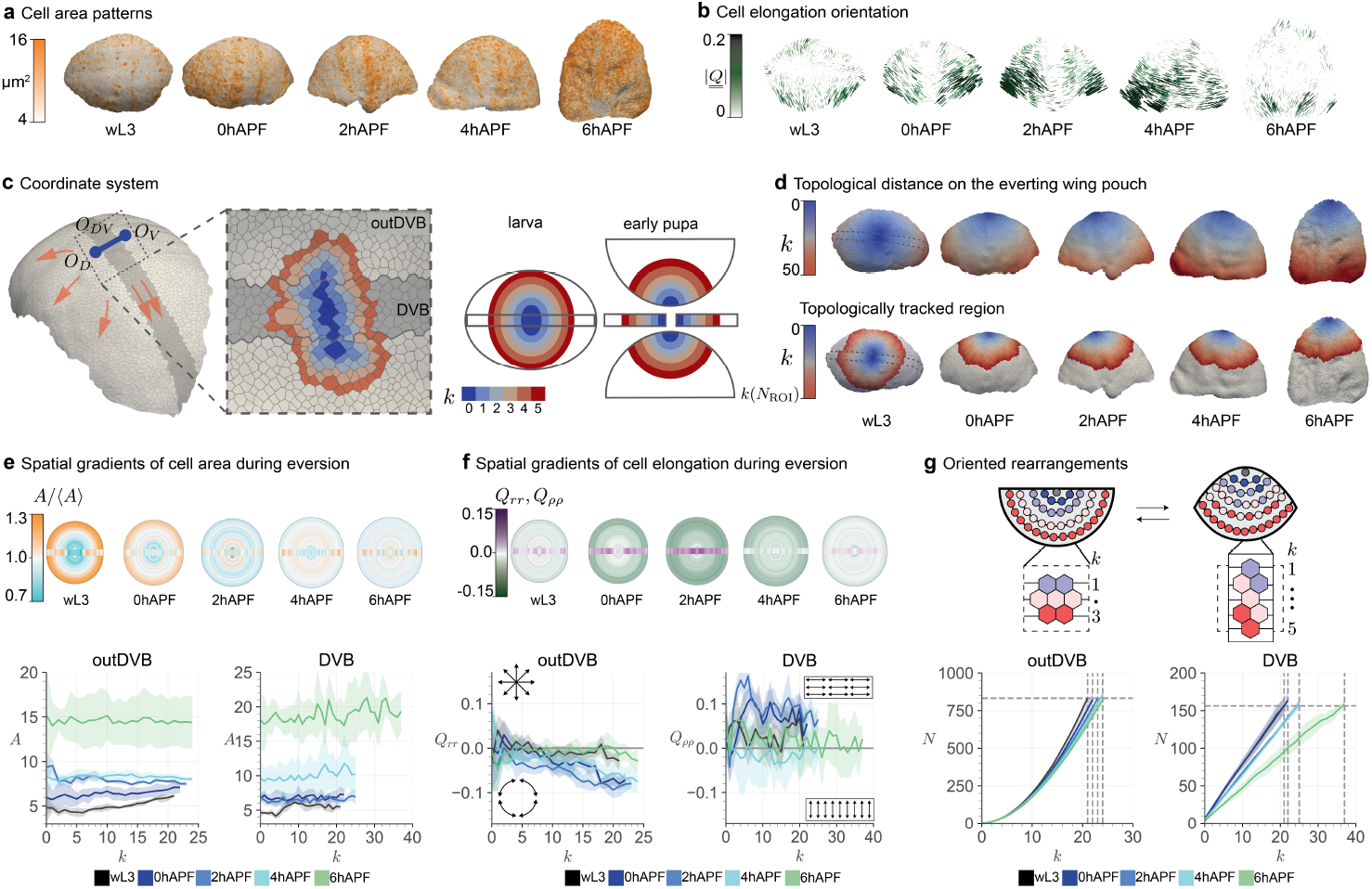
Topological tracking reveals spatial patterns of cell size changes, cell elongation changes, and cell rearrangements in the everting wing disc pouch: **a,b,d**, Cell measurements highlighted on the surface of representative examples of everting wing discs over time. At wL3, the full pouch is visible, whereas only the dorsal side is shown here for early pupal stages. **a**, Cells are colored by apical cell area. **b**, Bars highlight the orientation of locally averaged cell elongation 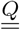 (projected onto 2D), and color indicates the elongation magnitude 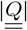 averaged over patches of size 350*µm*^2^. **c**, Segmentation of a wL3 pouch; the origins for the topological coordinates *k* (*O*_*D*_ and *O*_*V*_ in the outDVB region and a center line *O*_*DV*_ for the DVB) are highlighted in dark blue. The arrows indicate the direction of spatial coordinates that result from these origins, transversing along the DVB and radially for the outDVB. The inset shows the center region of the same wing pouch, where each cell up to *k* = 5 is colored by *k*. The origin cells are at *k* = 0 (dark blue). Note that the origin is a single cell for each side of the outDVB and a line of cells for the DVB. Due to the 3D nature of the wing pouch, the topological coordinate system is defined in one view for larval stages and in 4 separate imaging angles for early pupal stages (see schematic, right). **d**, The maximum *k* depends on the size of the segmented region (upper row). For the topologically tracked region, the maximum *k* may change due to rearrangements and is denoted *k*(*N*_ROI_) (lower row). **e-f**, Cell area (e) or cell elongation (f) spatially averaged over *k* (minimum five wing discs per stage). Dorsal and ventral are averaged together into ‘outDVB’. Geometric representations (top) show outDVB as half-circles and the DVB as a central rectangular box. **e**, Geometric representations highlight cell area gradients (*A/*⟨*A*⟩) within each stage. Lower panels show the cell area (*A*) as a function of *k* for all time points. **f**, The component of cell elongation (*Q*_*rr*_ for cells in the outDVB and *Q*_*ρρ*_ for cells in the DVB) is calculated relative to the origin for each cell of the respective region. This makes *Q*_*rr*_ the radial component of cell elongation, whereas *Q*_*ρρ*_ is effectively the cell elongation along the DVB (cartoon insets on the lower panel, see also Methods <u>7.8</u>). *Q*_*rr*_ and *Q*_*ρρ*_ are calculated as a function of *k*. In the upper panel, magnitudes are represented by color. **g**, Schematic (top) showing how we estimate cell rearrangements using topology. Each circle represents a cell in the outDVB region of the wing disc, colored by topological distance at the initial time point. If the number of cells per *k* decreases, the deformation by rearrangements is radial. Plots (bottom) show the number of cells in the wing disc pouch *N* contained within *k*. The horizontal line shows *N*_ROI_ for the wL3 stage; the vertical lines show corresponding (*k*(*N*_ROI_)) for each stage. In **e-g**, solid lines indicate the mean, and ribbons show 95% confidence of the mean.

To compare spatial patterns of cell behaviors over eversion time and across experiments, we define a coordinate system on the evolving 3D geometry. To this end, we use the cellular network topology to define the distance measure on the tissue surface. The topological distance between two cells is defined as the number of cells on the shortest path through the network from one cell to the other (See Extended Data Fig. S3a). We then use topological distance to define a coordinate system in the outDVB and DVB regions (Fig. 3c, Extended Data Fig.S3 and S4a,b, and Methods 7.8). The outDVB region consists of the dorsal and ventral halves, and we identify a single cell that defines the origin in each half (*O*_*D*_ and *O*_*V*_). In the DVB, we define the origin (*O*_*DV*_) as a line of cells transversing the DVB. The topological distance *k* to the origin defines a radial topological coordinate in each region (Fig. 3c,d).

During eversion, tissue previously hidden in the folds becomes visible. In order to compare cell behaviors at different time points, we need to identify a region of tissue that remains in the field of view throughout eversion. To this end, we count the number of cells *N*_ROI_ within the largest visible topological ring at wL3. The corresponding region of interest at later time points is then defined to be centered at the origin and containing the same number of cells. Since there are very few divisions and extrusions during eversion [14, 28], and because cells cannot flow across the DVB [34, 35], we expect that our regions of interest contain largely the same set of cells, and we refer to them as topologically tracked regions (Fig. 3d, Extended Data Fig. S5a,b).

Next, we quantify patterns of cell area (*A*) and radial cell elongation tensor 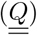 as a function of topological coordinate *k* throughout eversion (See Methods 7.7). We find that our topological coordinate system recapitulates previously reported gradients in cell area and radial cell elongation at earlier larval stages (Extended Data Fig. S4c,d). In outDVB at wL3, we observe a cell area gradient that relaxes gradually until 4hAPF (Fig. 3e). At the same time, cell elongation develops a gradient, with cells in the periphery elongating tangentially (Fig. 3f). Between 4h and 6hAPF, cells dramatically expand their area and tangential cell elongation completely relaxes. We do not observe gradients in cell area or cell elongation in the DVB. Instead, cell area expands while cell elongation along the DVB first increases up to 2hAPF and then decreases at 4hAPF (Fig. 3e,f).

Using topological distance allows us to extract spatial patterns of oriented cell rearrangements from snapshots of eversion. Radially oriented rearrangements lead to a decrease in the number of cells per *k*, whereas tangentially oriented rearrangements lead to an increase (see Fig. 3g). As a consequence, *k*(*N*_ROI_) changes based on the orientation and magnitude of rearrangements. We find that *k*(*N*_ROI_) increases with time (Fig. 3g), consistent with radially oriented cell rearrangements in outDVB and rearrangements oriented along the boundary in the DVB.

Together, these measured cell behaviors are a superposition of different radial patterns with the additional complexity of the DVB. Next, using our programmable spring model (Fig. 2), we ask how in-plane strains caused by these cell behaviors could drive 3D shape changes during eversion.

## 4 Active cell rearrangements drive tissue shape changes during wing pouch eversion

To be able to compare the output of the model to the 3D shape changes happening during eversion, we quantify the curvature and size dynamics of the apical surface of the wing pouch. We limit the analysis to the topologically tracked region and quantify the change in curvature from the wL3 stage along lines in the along-DVB and across-DVB directions (Fig. 4a, Extended Data Fig.S5, Methods 7.4). We focus on the stages between wL3 and 4hAPF, during which cell shape patterns have radial symmetry (Fig. 3a,b,e,f). We observe an overall curvature increase that is more pronounced in the across-DVB direction, peaking at the DVB, while flattening at the dorsal and ventral sides. Furthermore, the overall tissue area increases (Fig. 4a, Extended Data Fig. S5a).

**Fig. 4.**
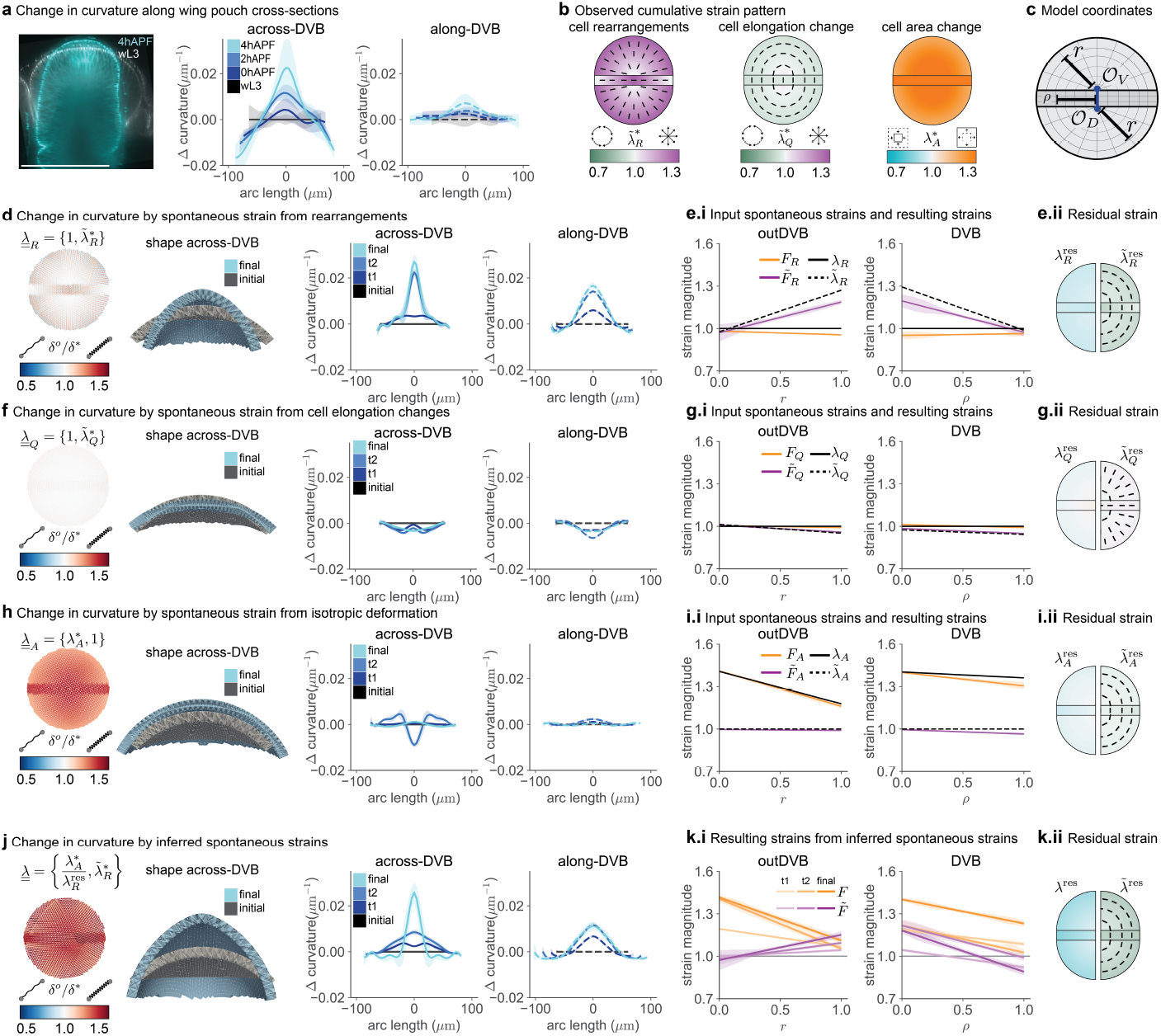
Inputting measured and inferred strains as spontaneous strain in the programmable spring model show that active rearrangements and cell area changes drive pouch morphogenesis: **a**, Overlay of a wL3 (white) and a 4hAPF (cyan) wing pouch (left) and plots of the average change in tissue curvature in the topologically tracked region in across-DVB (middle) and along-DVB (right) directions. **b,** Observed strain from cellular behaviors between time points wL3 to 4hAPF as a function of normalized distance from origin *r* and ρ. Observed strains arise from (left to right): rearrangements 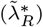, cell elongation changes 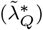, and cell area changes 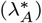. Half circles indicate the outDVB region; the rectangular box indicates the DVB. The color represents the magnitude of different strains; the bars visualize the direction of observed strain for 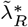 and 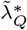. **c,** The model coordinates are designed to match the geometry of spatial patterns in the wing disc pouch (See also Fig. 3c). **d,f,h** Observed in-plane strain from rearrangements (**d,** 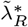), cell elongation changes (**f,** 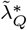), and cell area changes (**h,** 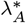) are inserted in the model as spontaneous strains by a change in rest length of the springs (δ°/δ^∗^). To compare the initial and final stages (corresponding to wL3 to 4hAPF), the model cross-section shows the shape in the across-DVB direction. The change in curvature of the model outcomes are plotted for all time points (right) in the across-DVB and along-DVB directions. The initial shape is a spherical cap with a radius resembling the wL3 stage. t1, t2, and final stages are the model results from the change in strains by 0, 2, and 4hAPF. 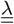 contains observed strains from the individual measured cell behaviors, while the other components (λ or 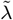) are set to 1. **e.i,g.i,i.i,** Input spontaneous strain for 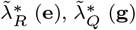, and 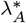 **(i)** at the final eversion time point (λ, 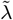) and the resulting strain that is achieved after relaxation of the model, which can be isotropic (F) and anisotropic 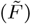. **e.ii,g.ii,i.ii**, Residual strain that remains at the final time point. The colors shows the magnitude of strain using the same range as indicated in Fig. 4b. Plots are split vertically to show the isotropic component (λ) on the left and the anisotropic component 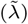 on the right. **j, k**, Model output and residual strains for the inferred spontaneous strains, following the same procedure as in **d-i**.

Next, we measure the strain field 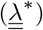 resulting from cell behaviors as a function of the distance from the origin, *r* or *ρ* (Fig. 4b, Extended Data Fig. S6, Extended Data Fig. S7, Methods 7.12). We quantify the isotropic component resulting from cell area changes 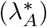 (Fig. 4b, Extended Data Fig. S7c). In the outDVB, we observe an area expansion 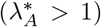 up to 2hAPF with a radially decreasing profile. In the DVB, we observe the buildup of a shallower gradient that is transiently paused from 0hAPF to 2hAPF. The contribution to the anisotropic component of 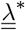 from changes in cell elongations 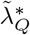 is small compared to the contribution by cell rearrangements 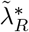 (Fig. 4b, Extended Data Fig. S7a,b,d, Methods 7.11). While 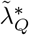 is tangential, following a shallow gradient, 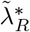 is radial and increasing with the distance from origin in the outDVB and decreasing in the DVB.

We next use the programmable spring model to test how the observed inplane cellular behaviors can cause tissue shape changes. We define the DVB and outDVB regions in the model, matching their relative sizes in the wing pouch (Fig. 4c, Extended Data Fig. S8). For each individual cell behavior and measured time point (wL3, 0hAPF, 2hAPF, and 4hAPF), we use the inplane 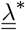 that we infer from each observed class of cell behaviors as examples of spontaneous strain 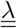 We use these to program the spring lengths in the model. For each insertion of spontaneous strain (model time points: initial, t1, t2, final, corresponding to the experimental time points), we relax the spring network quasi-statically to a force balanced state (Methods 7.9, 7.10). As the effective bending modulus of the wing disc is experimentally inaccessible, we fit the thickness of the model in an example scenario where all observed cell behaviors are input as spontaneous strains and use the same thickness thereafter (Extended Data Fig. S9a, Methods 7.14).

We first consider cell rearrangements as a possible source of spontaneous strain. When we only input 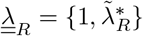 as spontaneous strain in the model, it alone creates a strong curvature increase, resembling many features of the data but without increasing tissue size (Fig. 4a,d). Note that 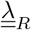 also introduces a difference in curvature change between the two directions, across-DVB and along-DVB, at the final stage.

After relaxing the spring network to a force balanced state, stresses due to residual strains remain. The stresses corresponding to these residual strains can drive passive responses in cell behaviors. The residual strains appear as a mismatch of spontaneous strains (input to the model, 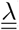) and strains resulting from changes in spring length during relaxation of the network, 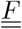 (Fig. 4e.i, Methods 7.10).

When we calculate the residual strains generated by spontaneous strain from rearrangements, we find that the anisotropic component of the residual strain 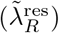 is tangentially oriented (Fig. 4e.ii, Extended Data Fig. S10d). This tangentially oriented strain is similar to the pattern of cell elongation changes 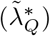 (compare Fig. 4b and 4e.ii), suggesting that these cell elongation changes are a passive response to spontaneous strain by rearrangements. To test this idea, we next consider cell elongation as possible source of spontaneous strain. When we only input 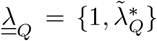 as spontaneous strain in the model, we observe that the spring network shape flattens at the center rather than curve, and cell elongations themselves do not lead to any further residual strains (Fig. 4f,g, Extended Data Fig. S10c). This result is consistent with cell elongation changes being a passive response to cell rearrangements and not driving tissue shape change during eversion.

Cell rearrangements as spontaneous strains also lead to residual isotropic compression (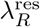, Fig. 4e.ii, Extended Data Fig. S10b). This residual could be compensated by spontaneous area change, which is also required by the observation that overall tissue size increases during eversion.

When we only input the isotropic strain, 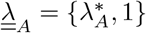 from observed cell area changes as spontaneous strain in the model, overall size increases with minimal curvature change (Fig. 4h). This result indicates that although cell area changes are an active behavior and lead to overall size increase, they do not significantly contribute to changes in tissue shape. However, there is a transient effect of cell area changes on tissue curvature at time point t2 (note dip in curve at t2 in Fig 4h). This transient effect in the scenario of spontaneous area strain only arises from the experimentally observed pause in cell area expansion in the DVB at 2hAPF as compared to the outDVB (Extended Data Fig. S7c, compare DVB and outDVB). This curvature difference disappears when the cell area in the DVB expands to match the outDVB at 4hAPF (Fig. 4h). Measuring 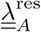, we find that cell area changes themselves create a small residual in the DVB (Fig. 4i, Extended Data Fig. S10). The anisotropic part of this residual could also contribute to the observed passive cell elongations.

Using these examples, we next infer the spontaneous strain patterns that drive tissue shape changes and govern cellular behaviors. We have found that both cell rearrangements and cell area changes are active and contribute to spontaneous strain. We therefore conclude that cell elongation is a passive elastic response and does not contribute to spontaneous strain. The total spontaneous strain, therefore, is composed of the anisotropic part of the observed strain due to rearrangements (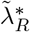, Fig. 4b) and the isotropic part of the observed cell area changes (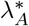, Fig. 4b) compensated by the isotropic part of the residual strain due to cell rearrangements (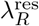, Fig. 4e.ii). When we input this total inferred spontaneous strain in the model, we find that we can account for the curvature and size changes observed in the everting wing pouch from wL3 to 4hAPF (compare Fig. 4j to Fig. 4a; see also Extended Data Fig. S9b, Supplemental movie 2). The patterns of residual strains generated by the model suggest that after eversion (at 4hAPF), cells experience elongation due to shear stress as well as area constriction due to compressive stresses (Fig. 4k, Extended Data Fig. S10f).

In summary, the good qualitative agreement between model output (Fig. 4j) and observed wing pouch curvature changes (Fig. 4a) indicates that the in-plane pattern of spontaneous strain by cell behaviors during eversion is sufficient to capture morphogenesis and that we have identified the most relevant active cellular events responsible for the pouch morphogenesis. Specifically, our data predict that altering cell rearrangements in the pouch should have a profound consequence for tissue shape change. We next test this prediction with a genetic perturbation.

## 5 Reduction of active cell rearrangements with MyoVI knockdown results in a tissue shape phenotype

Previous work in the wing disc pouch of earlier larval stages showed that cell rearrangements drive cell shape patterning [33]. This work suggested that patterns of active cell rearrangements self-organize via mechanosensitive feedback mediated by MyoVI. We therefore next investigate whether MyoVI knock-down in the wing pouch (Extended Data Fig. S11a) alters cell rearrangements during eversion and leads to a tissue shape phenotype.

Indeed, we observe that the MyoVI^RNAi^ wing disc pouch fails to form a flat bi-layer after eversion, even though its initial shape is similar to that of *wild type* (wt) (Fig. 5a.i). This phenotype is best captured in the behavior of curvature in the across-DVB direction (Fig. 5a.ii). Here, the curvature decreases in the center, in contrast to wt, where it increases. In the along-DVB direction, the curvature remains unchanged over time in the MyoVI^RNAi^ knockdown (Fig. 5a,b, Extended Data Fig. S11c,d).

**Fig. 5.**
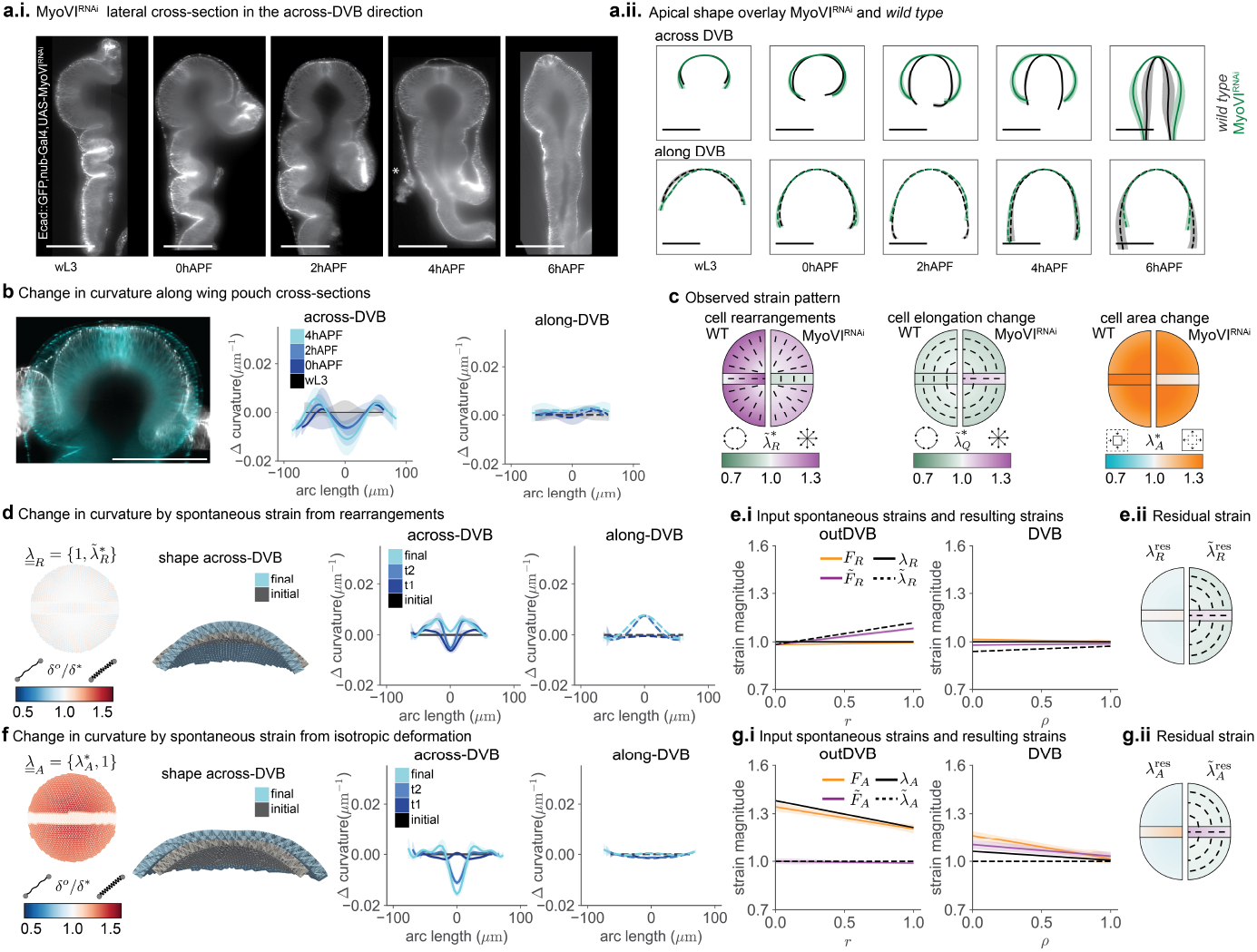
MyoVI^RNAi^ alters active cell behaviors and results in a tissue shape phenotype: **a**, MyoVI^RNAi^ phenotype during eversion (scale bars = 100 *μm*). Representative across-DVB cross-sections (**a.i**) and comparison of apical shape between MyoVI^RNAi^ and control (**a.ii**). **b,** Overlay of a wL3 (white) and a 4hAPF (cyan) MyoVI^RNAi^ wing pouch (left) and plots of the average change in tissue curvature in the topologically tracked region for across-DVB (middle) and along-DVB (right) directions. **c,** Observed strain from cellular behaviors in MyoVI^RNAi^ wing discs between time points wL3 to 4hAPF. Plots are split vertically with the observed strains for *wild type* (WT) for comparison on the left and MyoVI^RNAi^ on the right. Measured strains come from 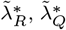, and 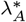. Quarter circles indicate the outDVB region, and the rectangular box indicates the DVB. The color represents the magnitude of different strains; the bars indicate the direction of observed strain for 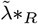 and 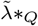. **d,f,** Observed in-plane behaviors are inserted in the model as spontaneous strains by a change in rest lengths of the springs (δ°/δ^∗^). The initial stage is a spherical cap with the radius taken to resemble the shape at the *wild type* wL3 stage. t1,t2, and final stages are the model results after a change in spring rest length according to observed strains from 0,2, and 4hAPF for MyoVI^RNAi^. 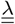 contains observed strains from rearrangements (**d**) or area changes (**f**), while the other components (*λ*_*R*_ or 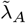) are set to 1. **e.i,g.i,** Input spontaneous strain ((λ and 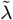) at the final eversion time point and comparison with the resulting strain that is achieved after relaxation of the model (*F* and 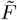). **e.ii,g.ii,** Residual strain that remains from the difference between input and resulting spontaneous strain at the final time point, plotted in the same way as Fig. 4e,g,i,k.ii.

Strikingly, other features of eversion, such as the opening of the folds and the removal of the peripodial membrane are unaffected by the MyoVI^RNAi^ knockdown (Fig. 5a, see 4hAPF), indicating that the cause for the altered shape is pouch-intrinsic. This result further supports the idea that tissue shape changes in the wing pouch during eversion are independent of other morphogenetic events happening in the wing disc and instead rely on active cell behaviors in the pouch.

Next, we quantify cell behaviors in MyoVI^RNAi^. While initially the gradients in cell areas and elongation are similar to wt (Extended Data Fig. S11e,f), the inferred strains from individual types of cell behaviors 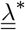 differ (Fig. 5c, Extended Data Fig. S12). From work in earlier larval stages, we expect oriented rearrangements to be reduced [33]. Indeed, we find that MyoVI^RNAi^ reduces the amount of radial cell rearrangements in the outDVB during eversion (Fig. 5c, Extended Data Fig. S11g). However, in the DVB, rearrangements are of opposite orientation as compared to wt eversion. Notably, we also see a complete lack of cell area expansion in the DVB (Fig. 5c, Extended Data Fig.S11e). The pattern of cell elongations in the outDVB is similar to wt, but in the DVB it is of perpendicular orientation (Fig. 5c, Extended Data Fig. S11f).

Using the programmable spring model, we test how the reduction of spontaneous strain due to cell rearrangements affects tissue shape changes. When we input 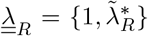 from cell rearrangements measured in MyoVI^RNAi^ as spontaneous strain in the model, we see only a slight increase in curvature in the final time point in both along- and across-DVB directions (Fig. 5d). Thus, we conclude that the reduction of cell rearrangements in MyoVI^RNAi^ as compared to wt contributes to the abnormal tissue shape changes happening during eversion in MyoVI^RNAi^. We find that the anisotropic component of the residual strain 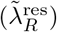 is small and tangentially oriented in the outDVB and radially in the DVB, similar to the cell elongation pattern (Fig. 5c,e, Extended Data Fig. S13d). If we input measured cell elongation changes as spontaneous strain in the model 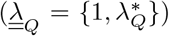, we do not recapitulate the observed tissue shape changes (Extended Data Fig. S14). This result suggests that the cell elongation changes in MyoVI^RNAi^ are a passive response to spontaneous strain by cell rearrangements, as in wt.

While the change in spontaneous strain due to rearrangements captures a significant portion of the difference between the wt and MyoVI^RNAi^ (compare Fig. 5b with Fig. 5d), it fails to recapitulate the finer progression of shape from wL3 to 4hAPF in MyoVI^RNAi^. In particular, the curvature at the final time point of the model calculation is not flattened in the center of the across-DVB direction, and the curvature increases slightly in both directions (Fig. 5b,d).

Thus, we proceed to input the observed cell area changes as spontaneous strains in the model 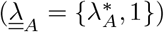. We find that they produce shape changes over time similar to those observed during MyoVI^RNAi^ eversion, recapitulating both the decrease in curvature in the center of the across-DVB direction and the lack of curvature change in the along-DVB direction (Fig. 5f, Supplemental movie 3). We conclude, therefore, that the subtle flattening in the pouch center in MyoVI^RNAi^ during eversion can be explained by the combination of cell area expansion in the outDVB with no area expansion in the DVB. This result highlights that, while cell area changes do not lead to a curvature change in wt, the difference in area expansion between the tissue regions results in the MyoVI^RNAi^ shape. In addition, although we do not observe cell area expansion in the DVB, the area expansion in the outDVB creates residual strains in both regions (Fig. 5g, Extended Data Fig. S13b). These residual strains have an anisotropic component that, together with the residual strains from cell rearrangements, account for the measured cell elongation patterns in MyoVI^RNAi^ (compare Fig. 5c,e.ii,g.ii)).

In sum, the results from the MyoVI^RNAi^ perturbation validate the idea that the wing disc pouch deforms like a shape-programmable material. First, by locally perturbing MyoVI, we show that we can alter the normal tissue shape change, even though the tissue outside behaves normally, demonstrating that the shape change is tissue autonomous. Second, we show that reducing the active cell rearrangements in the pouch significantly alters the tissue shape outcome, consistent with our theoretical model.

## 6 Discussion

In this work, we show that 3D epithelial tissue morphogenesis in the *Drosophila* wing disc pouch is based on in-plane spontaneous strains generated by active cellular behaviors. We develop a metric-free, topological method to quantify patterns of cell dynamics on arbitrarily shaped tissue surfaces, as well as a theoretical approach to tissue morphogenesis inspired by shape-programmable materials. These advancements together reveal the mechanics of tissue shape changes during wing disc eversion, showing that active rearrangements and active area expansion govern the 3D tissue shape and size changes.

We hypothesize that the organization of active behaviors during wing eversion arises from patterning during larval growth. First, the pre-patterned radial cell area gradient resolves during eversion, giving rise to a gradient of spontaneous strain in the outDVB. Second, the orientation of cell rearrangements follows that of earlier stages, indicating that the mechanosensitive feedback that was revealed in previous work is still active during eversion. Overall, this suggests a developmental mechanism through which mechanical cues at early stages organize cell behavior patterns that later resolve, resulting in a tissue shape change. Such behavior would resemble biochemical pre-patterning, in which cell fates are often defined long before differentiation.

Active, patterned rearrangements can robustly give rise to a specific target shape if the tissue is solid on the time scale of morphogenesis. Our work therefore reveals that the everting wing disc behaves as an elastic solid undergoing plastic deformation and demonstrates that the mere presence of rearrangements should not be taken as a sign of a fluid tissue with a vanishing elastic modulus. Many animal tissues with dynamic rearrangements could thus be in the solid regime and therefore be pre-patterned towards a target shape. Our work, inspired by shape-programmability of complex materials, reveals principles of shape generation that could be quite general. We therefore propose that many other morphogenetic events could and should be considered - and better understood - through the lens of shape-programmability.

## 7 Methods

### 7.1 Experimental model

All experiments were performed with publicly available *Drosophila melanogaster* lines. Flies were maintained at 25°C under 12hr light/dark cycle and fed with standard food containing cornmeal, yeast extract, soy flour, malt, agar, methyl 4-hydroxybenzoate, sugar beet syrup, and propionic acid. Adult flies were transferred to fresh food 2-3 times per week. Only males were studied for consistency and due to their smaller size. As *wild type*, we used the F1 offspring of a cross between *w-;ecad::GFP* and *w;nub-Gal4,ecad::GFP;;*.

### 7.2 *Drosophila melanogaster* lines

7.2 *Drosophila melanogaster* lines 437
| genotype | construct | origin |
| --- | --- | --- |
| *w-; ecad::GFP;;* | Ecadherin::GFP | Bloomington #60584 |
| *w; nub-Gal4, ecad::GFP;;* | nub-Gal4 | Bloomington #86108 |
| *w-; ecad::GFP, myoVI<sup>RNAi</sup>;;* | MyoVI<sup>RNAi</sup> | VDRC #37534 |

### 7.3 Image acquisition and processing

#### Sample preparation

Wing discs of larval stages were dissected in culture medium as previously described [36], without surface sterilization or antibiotics. Prepupal stages from 0 to 6 hAPF required a slightly different dissection strategy. Prepupae were marked by the time of white Pupa formation and collected with a wet brush after the required time interval. Next, pupae were placed on a wet tissue, cleaned with a wet brush to remove residual food, and transferred into glass staining blocks (see Ref. [36]) filled with dissection medium. To dissect the wing disc, a small cut was performed with fine surgical scissors (2.5 mm, FST 15000-08, Fine Science Tools GmbH) at the posterior end, which creates a small hole to release pressure. This allowed for the next cut to be performed at half the anterior-posterior length, separating the anterior and posterior halves. Next, the anterior part of the puparium was first opened at the anterior end by administering a cut just posterior to the spiracles and then a second cut was performed on the ventral side along the PD-axis. The pupal case was then held open with one forceps, and a second forceps in the other hand was used to remove the wing disc. To dissect wing discs from 0hAPF, the pupa is still soft enough to be turned inside-out after the cut that separates anterior and posterior halves, similar to larval stages.

#### Imaging

Imaging was performed with a Zeiss Lightsheet 1 system. Wing discs were mounted in capillaries (Zeiss, Capillary size 1, inner diameter ca. 0.68 mm) with 1 % low melting-point agarose (LMA, Serva, CAS 9012-36-6). LMA was prepared by mixing 1:1 of Grace’s insect medium and a 2% LMA stock solution in water. Wing discs were transferred into mounting medium in U-glass dishes, aspirated into the capillary at room temperature, and imaged immediately after the LMA solidified. The imaging chamber was filled with Grace’s insect medium (measured refractive index = 1.3424). For pupal stages, four imaging angles (dorsal, ventral, and 2 lateral) with 90° rotation were acquired; for larval stages, three imaging angles (dorsal and ventral in one, and 2 lateral) were acquired. Data from dual illumination was fused on the microscope using a mean fusion.

#### Multiview reconstruction

Multiview reconstruction was based on the BigStitcher plugin in Fiji [37, 38]. Images were acquired without fluorescent beads, and multiview reconstruction was done using a semi-automated approach. Individual views were manually pre-aligned. Thereafter, precise multiview alignment was computed based on bright spots in the data with an affine transformation model using the iterative Closest Point (ICP) algorithm. Next, images were oriented to show the apical side in XY and lateral in ZY. Lastly, images were deconvolved using point spread functions extracted from the bright spots and saved as tif files with a manually specified bounding box.

#### Surface extraction of 3D images for visualization

Surfaces shown in Fig. 1 and Supplemental movie were extracted from 2hAPF and wL3 images. To do so, we first trained a pixel classifier on the strong apical signal of Ecadherin-GFP of a different image of the same stage with napari-accelerated-pixel-and-object-classification [39, 40]. Feature sizes of 1-5 pixels were used to predict the foreground on the target image. Next, we used the pyclesperanto library [41] to select the largest labels and close gaps in the segmentation with the closing sphere algorithm. For additional gap-filling in the 2hAPF time point, we used vedo [42] to generate a pointcloud and extract the pointcloud density. When necessary, we applied some manual pruning of the segmentation in napari. We repeated this processing on the weak Ecadherin-GFP signal from the lateral membrane and subtracted the apical segmentation from the output. As a result, we achieved a full tissue segmentation that stops just below the apical junction layer. We then extracted the surface by the napari-process-points-and-surfaces [43] library and applied smoothing and filling holes. The visualizations were generated using Paraview [44]. Regions and directions of the cross-sections were annotated in Illustrator. Supplemental movies were created using paraview and Fiji [38].

### 7.4 Quantifying curvature of cross-sections

Tissue shape analysis was performed on multi-angle fused SPIM images. We used Fiji re-slicing tools to generate two orthogonal cross-sections along the apical-basal direction. Across-DVB is a cross-section along the center of the long axis of the wing disc. To find the center, we used the position of the sensory organ precursors and general morphology. The along-DVB cross-section follows the DVB and was identified by Ecadherin-GFP signal intensity. The apical pouch shape was outlined manually along both directions over the pouch region up to the HP-fold using custom Fiji macros. Subsequent pouch shape analysis was performed in Python. The tissue shape information was extracted form Fiji into Python using the Python ‘read-roi’ package.

The extracted apical shapes were aligned and rotated for each wing disc as follows. First, starting from the left-most point in the curve, we measure the arc length of the curve in the clockwise direction. The arc length of the *i*th point on the curve is given by

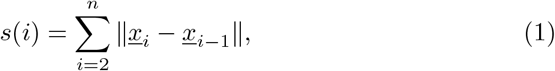

where *n* is the number of points in the discrete curve and *<u>x</u>*_*i*_ *=* (*x*_*i*_, *y*_*i*_) is the position vector of the *i*th point. We keep *s*(*i* = 1) = 0.

Next, we define the center of the curve at the middle and offset the arc lengths to have *s* = 0 at the center. This leads to negative arc lengths on the left side of the center and positive arc lengths on the right side of the center (Extended Data Fig. S1, S5).

In order to compute a mean curve from different wing discs of the same developmental stage, we translated and rotated the curves (Extended Data Fig. S1b). We translate each curve by setting their midpoints as the origin (0, 0). To rotate the curve, we compute the center of mass of the curve. Then, we define the new y axis as the line that joins the center of mass to the origin. Finally, for each curve, we smoothen and interpolate between the discrete points using spline interpolation. We use the *scipy*.*interpolate*.*UnivariateSpline* function of scipy [45]. To smoothen the spline, we define five knot points, one being the mid-point of the curve, and two others being at three-fourth and half of length from mid-point from either sides. Next, we compute the curvature of each curve using the following expression

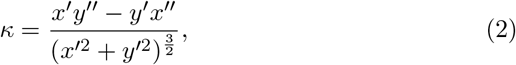

where ^*′*^ refers to the derivative with respect to the parameter of the curve, which is arc length in our case.

Finally, to compute an average curve, we get the average position vectors at arc lengths starting from a minimum arc length until a maximum arc length in intervals of 5*µm*. We do similar averaging for curvature values to get average curvature profiles.

To calculate the change in curvature, we normalized each curve from 0 to 1 and use a linear interpolation with 40 positions to subtract the initial from subsequent curvatures. We then re-introduced the average arc length for each each developmental stage for each of the normalized positions.

### 7.5 Segmentation of the apical junction network

To analyze cell shapes, we used four angles separated by 90° for the segmentation of early pupal stages, and a single angle for larval wing discs (Extended Data Fig. S2). Z-stacks from each imaging angle were denoised if necessary, by using the N2V algorithm [46], and the signal to background ratio was further improved by background subtraction tools in Fiji [38]. We made 2D projections of the Ecadherin-GFP signal in the Disc Proper layer as previously described [47]. Importantly, this algorithm also outputs a height-map image, which encodes the 3D information in the intensity of each pixel. The cells in the wing pouch were segmented using Tissue Analyzer and manually corrected [48]. We chose a bond length cutoff of 2 pixels (∼ 0.46*µm*). The ventral side for 0 hAPF was excluded from the analysis, as at this stage, the ventral region is never fully in view from any imaging angle. The number of wing discs per time point and the images for each region are indicated in Table 1. Images were rotated to orient distal down. Height-map images were rotated accordingly using imagemagickTM software (ImageMagick Development Team, 2021). We used Fiji macros included with TissueMiner [49] to manually specify regions of interest (ROIs). The DVB was identified based on Ecadherin-GFP signal intensity [50] and the dorsal vs. ventral pouch by their positions relative to global tissue morphology. For larval stages, the DVB, dorsal, and ventral regions were identified in one image. For images showing lateral views of pupal stages, the DVB was identified, whereas for images showing the outDVB region, the dorsal or ventral region and the cells next to the DVB were labelled. The cells next to the DVB were required as a landmark for topological analysis but were otherwise not analysed separately. We then ran the TissueMiner workflow to create a relational database.

**Table 1.**
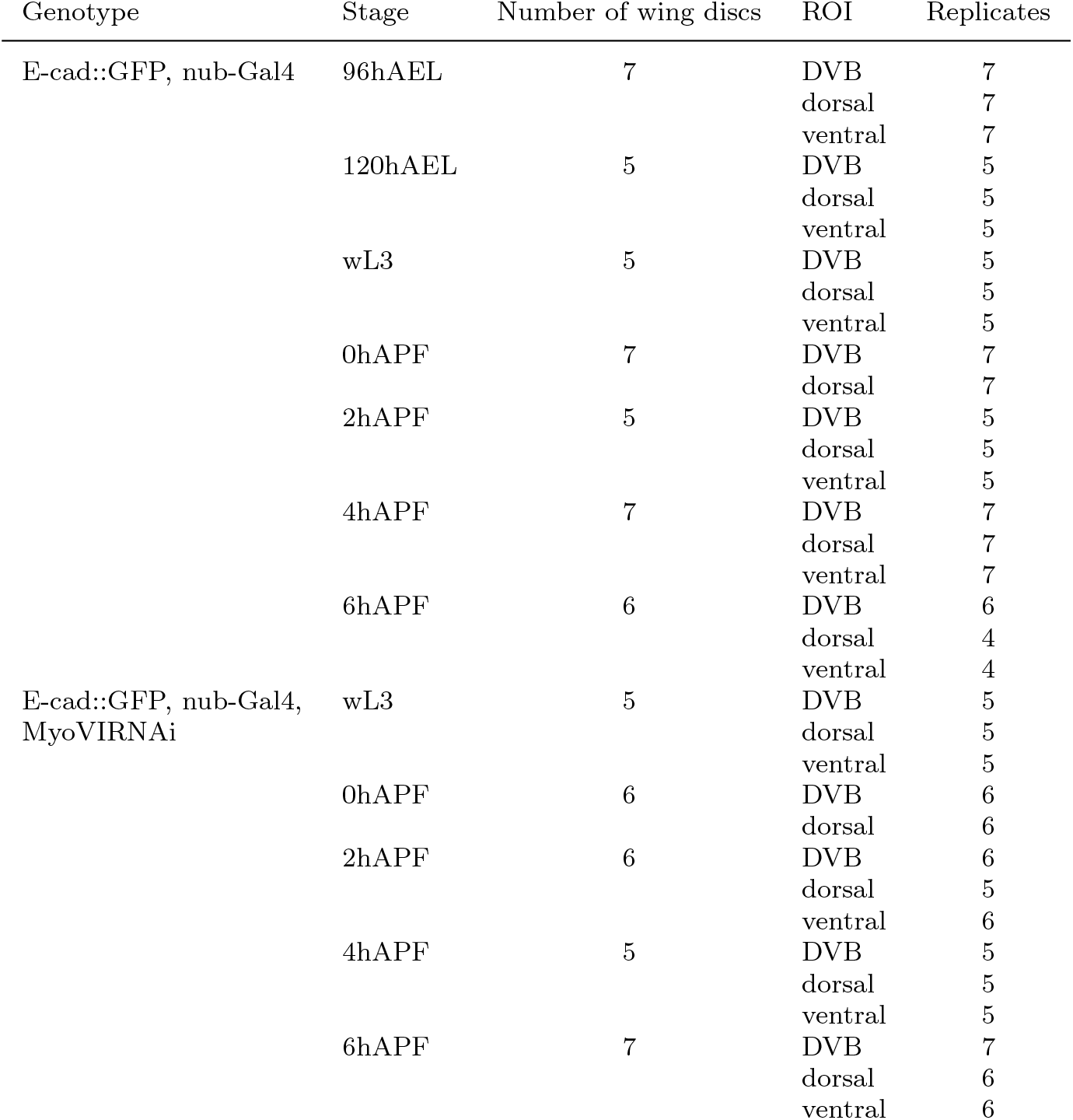
wing disc data.

### 7.6 3D cellular network

We represent the configuration of the cellular network by positions of the cell vertices, where three or more cell bonds meet, and their topological relations as in TissueMiner [49]. We extended TissueMiner to the third dimension using the information extracted from height-maps, as described in Methods 7.5.

### 7.7 Measurement of cell area and cell elongation tensor

Each cell *α* in the 3D network contains *N*^*α*^ vertices 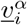, defining the network geometry. For every cell, we define a centroid *<u>R</u>*^*α*^, an area *A*^*α*^, and a unit normal vector 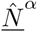 as

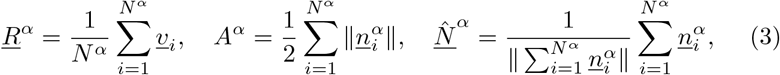

where 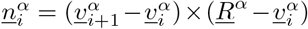 is the normal vector on the triangle formed by one edge of the cell and the vector pointing from the cell vertex to the cell centroid. It has a norm equal to twice the area of the triangle.

We then create a subcelluar triangulation by connecting the two consecutive vertices in every cell with its centroid {*<u>v</u>*_*i*_, *<u>v</u>*_*i*+1_*<u>R</u>*^*α*^}. This creates a complete triangulation that depends both on the vertex positions and the centroids of the cellular network.

Each triangle is defined by its three vertices { *<u>R</u>*_0_, *<u>R</u>*_1_, *<u>R</u>*_2_}, which define two triangle vectors *<u>E</u>*_1_, *<u>E</u>*_2_ and its unit normal vector <u>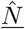</u>

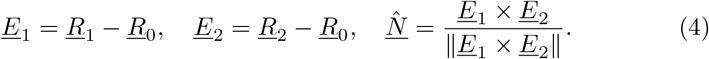

These vectors also define the local basis on the triangle. Using the triangle vectors, we can define the area of the triangle and the rotation angles *θ*_*x*_ and *θ*_*y*_ that rotate a vector parallel to the z-axis of the lab reference frame to the vector normal to the plane of the triangle

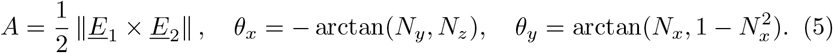

Here, arctan(*x, y*) is the element wised arc tangent of *x/y*, and *N*_*i*_ is a component of the unit vector normal to the triangle plane.

For each triangle, we define the triangle shape tensor 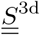 as a tensor that maps a reference equilateral triangle with area *A*_0_ lying in the xy-plane, defined by the vectors vectors *<u>C</u>*_*i*_to the current triangle

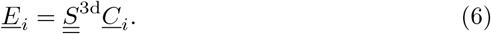

The vectors of the reference equilateral triangle are

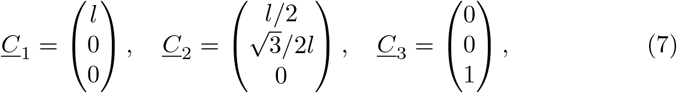

where the side length 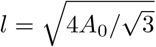 with *A*_0_ = 1.

The triangle shape tensor 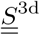 can be written in terms of a planar state tensor 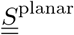 in the reference frame of the triangle as

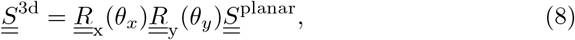

where 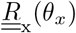 and 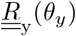 are rotations around the x and y axis, respectively. The angles *θ*_*x*_ and *θ*_*y*_ are defined in Eq. <u>5</u>. The planar triangle state tensor, represented by a 3x3 matrix with the *z* components set to 0, can be decomposed as

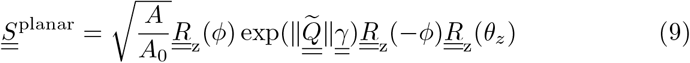

as in TissueMiner. Here, 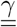 is a diagonal matrix with diagonal elements {1, −1, 0}, and 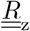 is the rotation matrix around the z-axis. *A* is the area of the triangle, 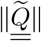 the magnitude of the elongation tensor, *ϕ* the direction of elongation in the xy-plane, and *θ*_*z*_ is the rotation angle around the z-axis relative to the reference unilateral triangle. The 3D elongation tensor 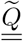 in the lab reference frame and the elongation tensor in the xy-plane of the triangle 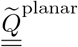 are related by

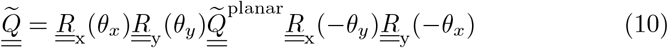

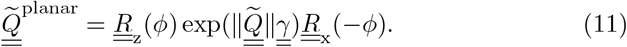

The magnitude of elongation is calculated as <u>[51]</u>

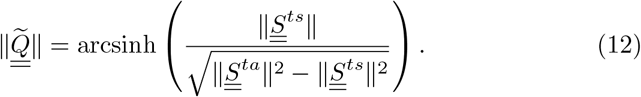

where 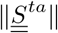 and 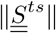 are the norms of the trace-antisymmetric and traceless-symmetric part of the planar triangle state tensor 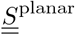, respectively. The angle of the elongation tensor is given by

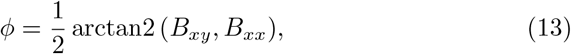

where *B*_*ij*_ are the components of the nematic part of the triangle state tensor 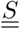 and arctan2(*x*1, *x*2) the inverse tangent of *x*1*/x*2, where the sign of *x*1 and *x*2 is taken into account. In this way, one can select the branch the multivalued inverse tangent function that corresponds to the angle defined by the point (*x*1, *x*2) in a plane.

We now define the cell elongation tensor as the area-weighted average of the corresponding triangle elongations

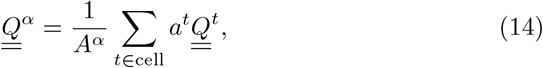

where *A*^*α*^ is the area of the cell, *a*^*t*^ the area of a triangle that overlaps with the cell, and 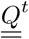 is the elongation tensor of that triangle.

To calculate the radial component of the cell elongation tensor relative to the origin in cell *α*, we first define the radial direction. To this end, we use a 3D vector *<u>r</u>* connecting the origin to the cell centroid and we project its direction 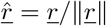 into the tangent plane of the cell, which defines the in-plane radial direction 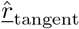. The tangent plane of the cell is defined by its normal vector 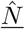 defined in Eq. <u>3</u>. We calculate the radial components of the cell elongation tensor as

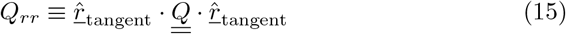

relative to the origin.

In the DVB, multiple cells form the origin. To calculate *Q*_*ρρ*_, the vector *<u>ρ</u>* connects the cell centroid to the averaged position of the topologically nearest cells of *k* = 0. We project its direction 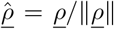 into the tangent plane of the cell *α*, which defines the in-plane direction 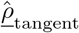 from DVB origin. We calculate the components of the cell elongation tensor as

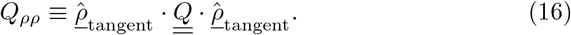

### 7.8 Topological distance coordinate system

To calculate topological distances between any two cells, we determine the topological network using the python-igraph library [52].

In each of the tissue regions, we define separate origins:

- outDVB region: To define the origin of the outDVB regions, we first determine the pouch margin cells as cells that live on the outermost row of the segmentation mask and do not overlap with the DVB ROI. Then, for each cell in the region, we calculate the shortest topological distance to the margin cells. This identifies the set of maximally distant cells that have the maximal shortest topological distance to the margin. The origin is then defined as the cell that is neighboring the DVB and is at the shortest metric distance to the averaged position of maximally distant cells. At larval stages, both dorsal and ventral sides of the outDVB region are visible, and an origin cell is defined on both sides.
- DVB region: We define the origin to consist of a line of cells transversing the DVB. At larval stages, the origin cells are defined as those cells within 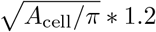 distance to a straight line connecting the dorsal and ventral center cells. For pupal stages, the origin cells for the DVB are defined as the first row of cells next to the margin of the segmentation mask on the distal side.

The so-identified origin cells serve as the origin for the topological distance (*k*) for each cell in the tissue. In this way, *k* follows the radial direction along the surface for the outDVB and the path along the the DVB for the DVB.

#### 3D visualization of cell properties

We visualize cellular properties and cell elongation tensors on the 3D segmentation mask using paraview [44].

To plot a rank 2 tensor, like the cell elongation tensor, we take the largest eigenvalue of 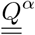 as the norm of elongation and the corresponding eigenvector as the direction of elongation that we can plot to the surface. Note that for cells / patches that are reasonably flat, the eigenvector with the eigenvalue closest to zero is (almost) parallel to the normal vector on the patch.

#### Spatial analysis of cell properties

We acquired data for 5 to 7 wing discs of each developmental stage. Images that were not of segment-able quality were excluded from the analysis. We average cell properties by *k* between dorsal and ventral for the outDVB and between images from both sides of the DVB. We used a cell area-weighted average for elongation. The 95 % confidence interval and the statistic mean for each developmental stage is calculated via bootstrap re-sampling with 10.000 repeats.

### 7.9 Mechanics of the programmable spring lattice

We use a programmable spring lattice in the shape of a spherical cap to model the wing disc pouch, which is an epithelial monolayer.

#### Approximating the wing disc pouch as a spherical cap

We calculate the average radius of curvature of the apical side of the wing disc pouch at wL3 stage in the topologically tracked region as *R* = 77.66*µm*. The angular size of the spherical cap, denoted by *θ*_*M*_, is given by

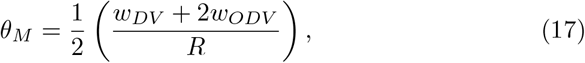

where *w*_*DV*_ is the width of the DVB and *w*_*ODV*_ is the average in-surface distance from the DVB to the periphery of the outDVB region (Extended Data Fig. S8a). We calculate *w*_*DV*_ = 15*µm* and *w*_*ODV*_ = 59.77*µm*. Using these calculated dimensions, we determine *θ*_*M*_ = 49.63^*°*^.

#### Generating the lattice

We first generate a triangular lattice in the shape of a hollow sphere, keeping the radius of curvature *R* calculated above. This lattice was obtained using the function *meshzoo*.*icosa sphere* available in the Python package Meshzoo (www.github.com/meshpro/meshzoo). In this function, we set the argument *refine factor* = 30, which leads to edges of length 3.11 ± 0.18*µm*. This edge length was found to be small enough to prevent computational errors in the simulations of this study. We then cropped the spherical lattice to obtain a spherical cap of angular size *θ*_*M*_ (calculated above, Extended Data Fig. S8b). Next, we place a second layer at the bottom of this lattice at a separation of *h*. This new layer is identical to the original lattice in terms of the topology of the lattice network but is rescaled to have a radius of curvature of *R* − *h*. We connect the two layers with programmable springs using the topology shown in the inset of Extended Data Fig. S8c. The lattice obtained this way represents an elastic surface of thickness *h*, which can be changed to tune the bending rigidity of the model. Vertices typically have 13 neighbors (6 on their own layer and 7 on the other layer). However, six to eight vertices out of about 3220 vertices in the whole network form point defects. These vertices have 11 neighbors.

In order to remove any possible effects coming from the lattice structure (angle of edges or degree of connectivity), we perform simulations for each condition by taking spherical caps from 50 different regions of the sphere and averaging the result. We see only very small variability in the final shape, quantified by the standard deviation of the curvature change profiles in our model results. Thus, we conclude that the lattice structure does not affect our results.

#### Elastic energy of model

The edges of the lattice act as overdamped elastic springs with rest lengths equal to their initial lengths. Hence, the model is stress-free at *T* = 0

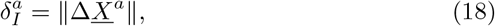

where *a* denotes a single spring; Δ*<u>X</u>*^*a*^ denotes the spring vector given by *<u>X</u>*^*β*^ − *<u>X</u>*^*α*^, where *α* and *β* are the vertices at the two ends of spring *a* and *<u>X</u>*^*α*^ denotes the position vector of vertex *α*. During a consequent time step *T*, the rest length of spring *a* 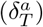 can differ from its current length *δ*. The elastic energy of this state for the whole lattice is given by

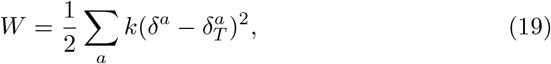

where the sum is over all springs of the network and *k* represents the spring constant. At each computational time step *T*, the model tries to find a preferred configuration by minimizing W, hence *T* acts as a “quasi-static time step”. To minimize the energy of the model at a given *T*, we use overdamped dynamics with smaller time steps *τ*, which restart for each new quasi-static time step *T*.

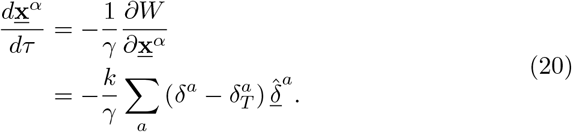

Here, *γ* represents the friction coefficient. **<u>x</u>**^*α*^ corresponds to the current position of the vertex *α. δ*^*a*^ is the length of the springs connected to vertex *α*. 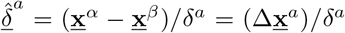 represents the unit vector along the spring *a* that connects vertices *α* and *β*.

We relax the model at each quasi-static time step *T* to achieve force balance by updating the positions of the particles using

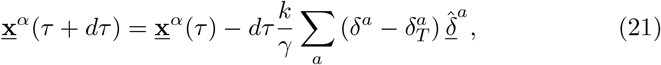

where 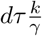 was set to 0.01 (ensuring no numerical artifacts).

The particles were moved until the average movement of the particles ⟨ ∥ **<u>x</u>**^*α*^ (*τ* + *dτ*) − **<u>x</u>**^*α*^ (*τ*)∥⟩*/R* reduced to 10 ^*−*9^, where *R* is the radius of curvature of the outer surface of the spherical cap in the initial stress-free state.

### 7.10 Spontaneous strain tensor

Tissue shape change during development is modelled in this work as the appearance of spontaneous strains, a change in the ground state of local length scales. This notion can be captured with a spontaneous strain tensor field, 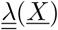, a rank 2 tensor. Each component corresponds to the multiplicative factor by which the rest lengths of the material changes in a particular direction. In some general coordinate system, we can write 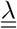 as

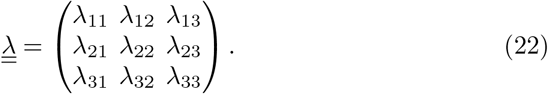

We choose the coordinate system so that it aligns with our desired deformation pattern. In this case, 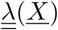 is in a diagonal representation :

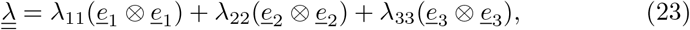

where the basis vectors are chosen such that *<u>e</u>*_1_, *<u>e</u>*_2_ are surface tangents while *<u>e</u>*_3_ is surface normal. In general, we keep *λ*_33_ = 1, since we do not input any spontaneous strains along the thickness of the model.

The surface components of 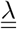 can be further broken down into isotropic and anisotropic components. Isotropic deformation changes the local area of the surface by changing the local lengths equally in all directions. Anisotropic deformation increases the local length in one direction while decreasing the local length in the other direction so as to preserve the local area. Thus, we decompose the deformation as a product of isotropic and anisotropic contributions.

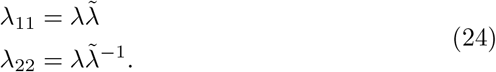

Then, the spontaneous deformation tensor can be written as

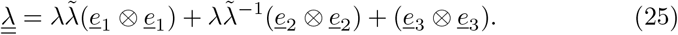

Finally, as 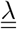 is a field, each of the components in the above equation generally depend on the location on the surface, *<u>X</u>*.

#### Discretizing 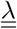

As our spring lattice is discrete in nature, we use the following strategy to discretize 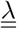. For a single spring, 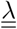 is an average of the value of 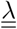 on the two ends of the springs.

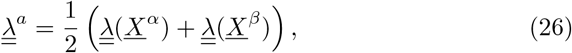

where *α* and *β* are the two vertices of the spring *a*.

#### Assigning new rest lengths to springs

The initial length of spring *a* connecting vertices *α* and *β* is given by

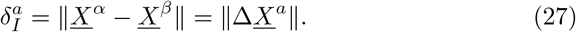

To assign new rest lengths, we use

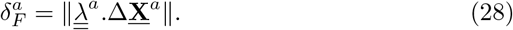

Note that we assign a new rest length to any spring *a* based on the positions of its vertices (*<u>X</u>*^*α*^ and *<u>X</u>*^*β*^), independent of the layer in which these vertices lie (top and bottom).

#### Implementing shape change over time

We increase the spontaneous strain slowly to model the slow build up of stresses due to cell behaviours. Hence, we first calculate the target rest length of springs 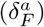. At each time step, we assign a rest length 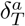 and minimize the energy of the model. We increase 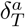 in a simple linear manner from 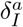 to 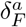

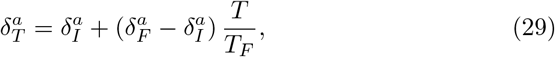

where *T*_*F*_ is the number of quasi-static time steps in which the whole simulation takes place. Note that within each time step, the lattice is brought to a force balance state. The simulations were performed for different choices of *T*_*F*_ (1, 2, 5), but we found that the differences in output shapes were undetectable. Still, *T*_*F*_ = 5 was chosen to simulate the slow appearance of spontaneous strains.

#### Measuring resulting strains in model

In our spring model, displacements are defined by positions of vertices and we define the deformation gradient tensor 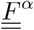 at each vertex *α* of the network.

For each spring *a* emerging from the vertex *α*, the deformation gradient tensor should satisfy

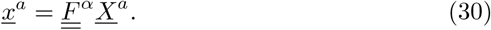

However, 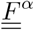 contains 9 degrees of freedom, while there are 13 springs for each vertex and therefore 13 independent equations to be satisfied. Note that six to eight vertices out of about 3220 vertices in the whole network form point defects and thus have 11 springs. Therefore, we define 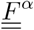 as the tensor that L best satisfies conditions in Eqs. <u>30</u> by minimizing the sum of residuals squared

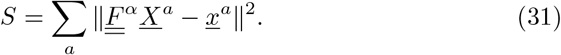

This is an ordinary least squares (OLS) problem split into three independent basis vectors. We solved this OLS using the Numpy method *numpy*.*linalg*.*lstsq* in cartesian coordinates [<u>53</u>]. We then express 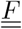 in the coordinate system corresponding to vertex *α* in the model explained above. From this, we calculate the isotropic (*F*) and anisotropic 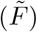 components using

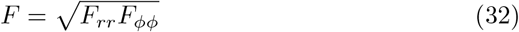

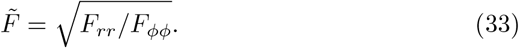

Finally, we compute 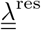as

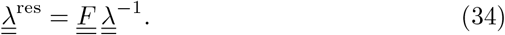

The isotropic (*λ*^res^) and anisotropic components 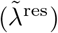 of 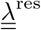 are calculated in the same way as for 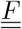.

### 7.11 Nematic director pattern on spherical surface

In the initial state of the model, we specify a coordinate system on the spherical surface in different regions (outDVB and DVB). These coordinate systems are chosen such that the observed nematic patterns of spontaneous strains 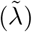 align with the major axes of the chosen coordinate systems.

We first define the origins in our model similar to the origins defined in the data (Extended Data Fig. S8). To do so, we first measure *θ*_*D*_*V*. The coordinates of *O*_*D*_ and *O*_*V*_ are then given by (±*R* sin(*θ*_*DV*_ */*2), 0, *R* cos(*θ*_*DV*_ */*2)) in the cartesian coordinate system. The center for the DVB region is given by the line *O*_*DV*_ which joins *O*_*D*_ and *O*_*V*_.

In the outDVB region, we have a coordinate system in which the basis vectors are given by *<u>e</u>*_*r*_, *<u>e</u>*_*ϕ*_, *<u>e</u>*_*h*_ (Extended Data Fig. S8c). *<u>e</u>*_*h*_ is simply the normal vector on the spherical surface. To calculate *<u>e</u>*_*r*_ at a point, we draw a vector from the origin in this region (*O*_*D*_ or *O*_*V*_) to the point. We then take a projection of this vector onto the tangent plane of the surface and normalize it to give us a unit vector. In this way, we calculate *<u>e</u>*_*r*_ as a surface tangent vector emanating radially outwards from the origins of the outDVB regions. *<u>e</u>*_*ϕ*_ is then the direction perpendicular to *<u>e</u>*_*r*_ and *<u>e</u>*_*h*_. For each point in the outDVB region, we calculate the geodesic distance between the point and the center point of its region. We then normalize this distance by the maximum geodesic distance from the center calculated in this region. This gives us a scalar coordinate *r* which varies from 0 to 1.

In the DVB region, the basis vectors are given by (*<u>e</u>*_*ρ*_, *<u>e</u>*_*w*_, *<u>e</u>*_*h*_) (Extended Data Fig. S8c). *<u>e</u>*_*h*_ is simply the normal vector on the spherical surface. To calculate *<u>e</u>*_*ρ*_ at a point, we draw a vector from the nearest point on *O*_*DV*_ to the point. We then take a projection of this vector onto the tangent plane of the surface and normalize it to give us a unit vector. In this way, we calculate *<u>e</u>*_*ρ*_ as a surface tangent vector emanating outwards from the center line of the DVB region as well as parallel to the DVB. *<u>e</u>*_*w*_ is perpendicular to *<u>e</u>*_*ρ*_ and *<u>e</u>*_*h*_. For each point in the outDVB region, we calculate the shortest distance between the point and the center line of the DVB region. We then normalize this distance by the maximum distance from the center line in the DVB region. This gives us a scalar coordinate *ρ*.

For the simple examples presented in Fig 2c (except Fig 2c.iii), *θ*_*DV*_ was set to be 0 to have a simple radial coordinate system. For Fig 2c.iii, *θ*_*DV*_ *> θ*_*M*_.

### 7.12 Extracting the strain pattern from segmented images

To quantify the strain due to different cell behaviors along the basis vectors of the chosen coordinate system, we compare cells within topologically tracked bins between two different developmental stages.

#### Tracking location between developmental stages

We leverage the topological distance coordinate system to track locations between discs. Each topological ring *k* is given a value *N* which denotes the cumulative number of cells from the topological origin defined in each region (*O*_*D*_, *O*_*V*_, and *O*_*DV*_). We use *N* to track the location in our static images of different discs at different developmental stages.

#### Observed strain due to cell area change

Cell area scales with square of the distance between cell vertices. Thus, the factor by which the local lengths change in all directions is given by

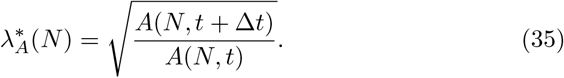

Here, *t* corresponds to an initial developmental stage, and *t* + Δ*t* corresponds to a later developmental stage. *A* refers to the average cell area evaluated at *N*.

#### Observed strain due to cell elongation change

Each cell is given a cell elongation tensor 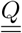 that is the average of further subdivisions of the cell polygon into triangles (Methods 7.7). Each triangle can be circumscribed by an ellipse, the centroid of which coincides with the centroid of the triangle. According to [54], the length of the long axis of the ellipse is given by 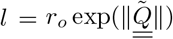, where *r*_*o*_ is the radius of a reference equilateral triangle. The length of the short axis of the ellipse is given by 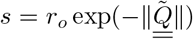. The axes of the ellipse match with the radial and tangential directions if the off-diagonal components *Q*_*rϕ*_ or *Q*_*ρϕ*_ are approximately 0. This was the case for our data as well. The length scale associated with the radial direction is *l* if *Q*_*rr*_ or *Q*_*ρρ*_ is positive and *s* if *Q*_*rr*_ or *Q*_*ρρ*_ is negative. Thus, we get a measure of the length scales along the radial direction, which we denote by *L* and is given by

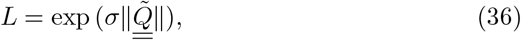

where *σ* is the sign of *Q*_*rr*_ or *Q*_*ρρ*_.

We then average *L* within each ring and compute a ratio of the length scales along the radial direction between two developmental stages by computing

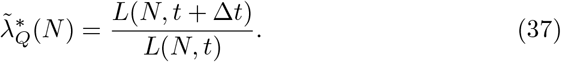

#### Observed strain due to cell rearrangements

Rearrangements lead to anisotropic deformation of the tissue. In our topological coordinate system, radially oriented rearrangements lead to an increase in the number of rings needed to accommodate some fixed number of cells (Extended Data Fig. S6). Similarly, tangential rearrangements would lead to a decrease in the number of topological rings. Thus, by measuring the change in the number of rings needed to accommodate some fixed number of cells, we can estimate the deformation due to the net effect of radial and tangential rearrangements.

In a tissue region at developmental stage *t*, let us consider a single ring with index *k* and cumulative number of cells *N*. Ring *k* contains Δ*N* cells given by *N* (*k, t*) − *N* (*k* −1, *t*). By construction, the number of rings needed to contain Δ*N* cells at location *N* is given by *n*(*N, t*) = 1. For a later developmental stage, *t* + Δ*t*, we estimate *n*(*N, t* + Δ*t*) which is the number of rings that contain Δ*N* cells at the location *N*. This is done by taking the difference between *k* values evaluated at *t* + Δ*t* and at locations *N* (*k*− 1, *t*) and *N* (*k, t*) (see also Extended Data Fig. S6)

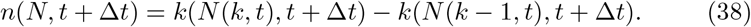

As *n*(*N, t*) and *n*(*N, t* + Δ*t*) are measures of the number of topological rings, they represent the radial topological length scales that change due to cell rearrangements. Thus, the strain due to cell rearrangements is quantified by

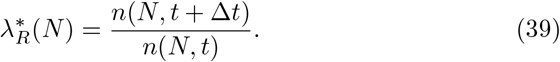

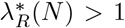 represents radial extension of the tissue due to radially oriented rearrangements, while 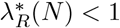 represents tangential extension.

#### Observed strain due to combination of cell elongation change and cell rearrangements

The combined strain due to cell elongation change and cell rearrangements is given by

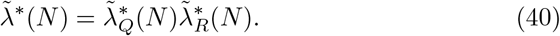

#### Mapping locations between wing disc images and model

In the model, we have a dimensionless scalar coordinate in the outDVB and DVB regions varying from 0 to 1. In the data as well, we prescribe a scalar coordinate to each topological bin. To do so, we calculate the path length in *µm* of the shortest path along cell centers from each cell to the origin and average this path length for each topological bin. For each bin, we normalize this path length by the average path length of the outermost topological bin in the corresponding region (DVB or outDVB). We call this normalized path length *r* for the outDVB region and *ρ* for the DVB region. Due to our normalization, *r* and *ρ* run from 0 to 1, similar to the model.

Thus, we are able to map any topological ring (identified by *k* and *N*) to a scalar coordinate *r* in outDVB or *ρ* in DVB. In Methods <u>7.11</u>, we explain the mapping between *r* and *ρ* to the cartesian coordinates of the vertices in the model given by *<u>X</u>*^*α*^, where *α* is a vertex. Using this mapping, any strain component, for example 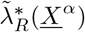, on the model vertex *α* can be evaluated from a corresponding 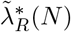

### 7.13 Quantifying curvature of model cross-sections

To quantify the curvature of the model output, we first isolate the top layer of the lattice. Then, we take the along-DVB cross-section (XZ plane) and the across-DVB cross-section (YZ plane). To take the cross-section, we record the points of intersection of the in-surface springs with the respective plane of the cross-section. From this, we get a discrete set of points that are ordered along their horizontal position to get a counter-clockwise curve. This curve data is now similar to the data we obtain from segmented images. Hence, we apply the exact same procedure described above to quantify the curvature of the model output.

### 7.14 Tuning thickness

We tune the thickness of the model in order to change the bending modulus. We first perform simulations by inputting all cell behaviours (cell area changes, cell elongation changes and cell rearrangements) combined as spontaneous strains. We perform this simulation for different thicknesses *h/R* = 0.05, 0.1, 0.15, where *h* is the thickness of the model and *R* is the radius of curvature of the top surface of the initial state of the model (Extended Data Fig. S9a). We find that *h/R* = 0.1 gives us the best matching of the curvature change profiles with the wing disc pouch. Fixing *h/R* = 0.1, we perform further analysis to infer the spontaneous strains in the wing disc pouch. Inputting these inferred spontaneous strains, we again performsimulations for different thicknesses. We find *h/R* = 0.1 still matches the wing disc pouch curvature change values best (Extended Data Fig. S9b).

### 7.15 Data availability

Imaging and model data are available upon request without restriction.

### 7.16 Code availability

Uncited code is available upon request without restriction.

## Acknowledgements

We thank Christian Dahmann, Anne Classen, and past/present members of the Eaton, Dye, Jülicher, Modes, and Popović teams for discussions on the project prior to publication. We also thank Bruno C. Vellutini for sharing his expertise on the multi-angle reconstruction of light sheet data. Miki Ebisuya, Stephan Grill, and Pavel Tomančák gave thoughtful feedback on the manuscript. We also thank the Light Microscopy Facility, the Computer Department, and the Fly Keepers of the MPI-CBG, as well as the Bioimage Analysis group at PoL for their support and expertise. This work was funded by Germany’s Excellence Strategy - EXC-2068 - 390729961- Cluster of Excellence Physics of Life of TU Dresden, as well as grants awarded to SE from the Deutsche Forschungsge-meinschaft (SPP1782, EA4/10-2) and core funding of the Max-Planck Society. NAD additionally acknowledges funding from the Deutsche Krebshilfe (MSNZ P2 Dresden). AK was funded through the Elbe PhD program. CD acknowledges the support of a postdoctoral fellowship from the LabEx “Who Am I?” (ANR-11-LABX-0071) and the Université Paris Cité IdEx (ANR-18-IDEX-0001) funded by the French Government through its “Investments for the Future”. We dedicate this work to our coauthor Prof. Dr. Suzanne Eaton, who tragically passed away near the beginning of the project.

## Author Contributions

JFF: Conceptualization, Methodology, Software, Validation, Formal analysis, Investigation, Data Curation, Writing - Original Draft, Visualization

AK: Conceptualization, Methodology, Software, Validation, Formal analysis, Investigation, Data Curation, Writing - Original Draft, Visualization

JP: Methodology, Software, Validation, Data Curation

CD: Methodology, Validation

SE: Conceptualization, Resources, Funding acquisition, Supervision

MP: Methodology, Validation, Writing - Original Draft

FJ: Conceptualization, Writing - Original Draft, Supervision

CDM: Conceptualization, Methodology, Resources, Writing - Original Draft, Project administration, Supervision, Funding acquisition

NAD: Conceptualization, Methodology, Resources, Writing - Original Draft, Project administration, Supervision, Funding acquisition

