## Extended Data Figures for "Active shape programming drives *Drosophila* wing disc eversion"

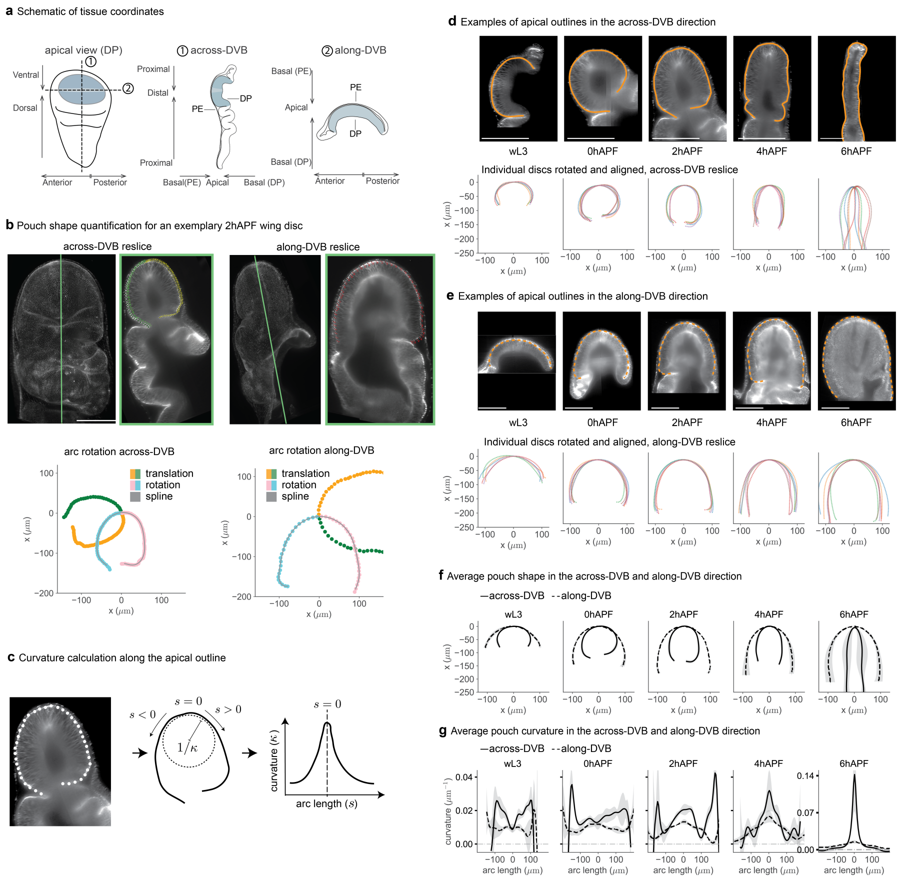


**Fig. S1 Pouch shape quantification**: **a**, Schematic of a wL3 wing disc, top down and in the two cross-sections. **b**, Examples of an across-DVB and along-DVB re-slice in a 2hAPF wing disc (maximum projection left, re-slice right). Wing discs are oriented to have Anterior-Posterior along the horizontal axis, Proximal-Distal (and Dorsal-Ventral for larval stages) along the vertical axis, and Apical-Basal in Z (and Dorsal-Ventral for early pupal stages). The across-DVB direction is identified on a maximum projection along Z and goes along the center of the long axis of the wing disc; the along-DVB direction is identified from the across-DVB direction and goes along the DVB. The apical arc is generated by spline interpolation from manually annotated points along the apical outline, then translated and rotated. The center of the arc is the position of the DVB for the across-DVB direction, and the mid-point for the along-DVB direction. **c**, Schematic explaining the curvature quantification: We extract the apical surface from image cross-sections (left); The curvature ($\kappa$) is calculated along the apical outline ('arc'); The arc is centered around the mid point ($s = 0$) of the curve and the arc length ($s$) is calculated along the curve in both directions from the mid point (middle, see also Methods 7.4 and Equation 2).**d,e**, Exemplary images of the wing pouch of each stage in the across-DVB (**d**) and along-DVB cross-section (**e**). Scatter plots below show the aligned and rotated outlines for all wing discs of each stage. **f**, Average shape shape and standard deviation (ribbon) over a minimum five discs for all stages. **g**, Mean curvature (solid line) and standard deviation (ribbon) over arc length ($\mu m$) The y-axis scale is the same through wL4 to 4hAPF and separately indicated for 6hAPF. **f,g**, Anterior is left in the along-DVB direction, and dorsal is left for the across-DVB direction. Scale bars = $100\mu m$.

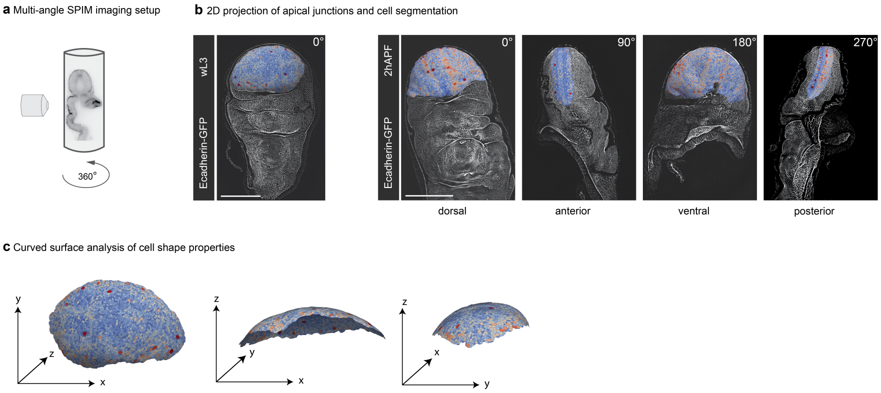


**Fig. S2 Multi-angle segmentation strategy and reintroduction of the 3D information to the cell segmentation mask**: **a**, We image wing discs from different angles using an agarose tube mounted in a multi-angle light sheet microscope. **b**, A single angle suffices to segment the pouch in larval wing discs, whereas 4 angles with 90° difference are required for the segmentation of all pouch regions in early pupal stages. For each angle, the apical surface projection is shown with an overlay of the segmentation in the pouch, where cells are colored by their apical area. **c**, Different angles of a wL3 stage segmentation; visualizations generated in Paraview. Using the height-map generated by the apical surface projection, we find the 3D shape of the tissue surface and calculate cell properties on this curved surface. Scale bars = 100 $\mu m$.

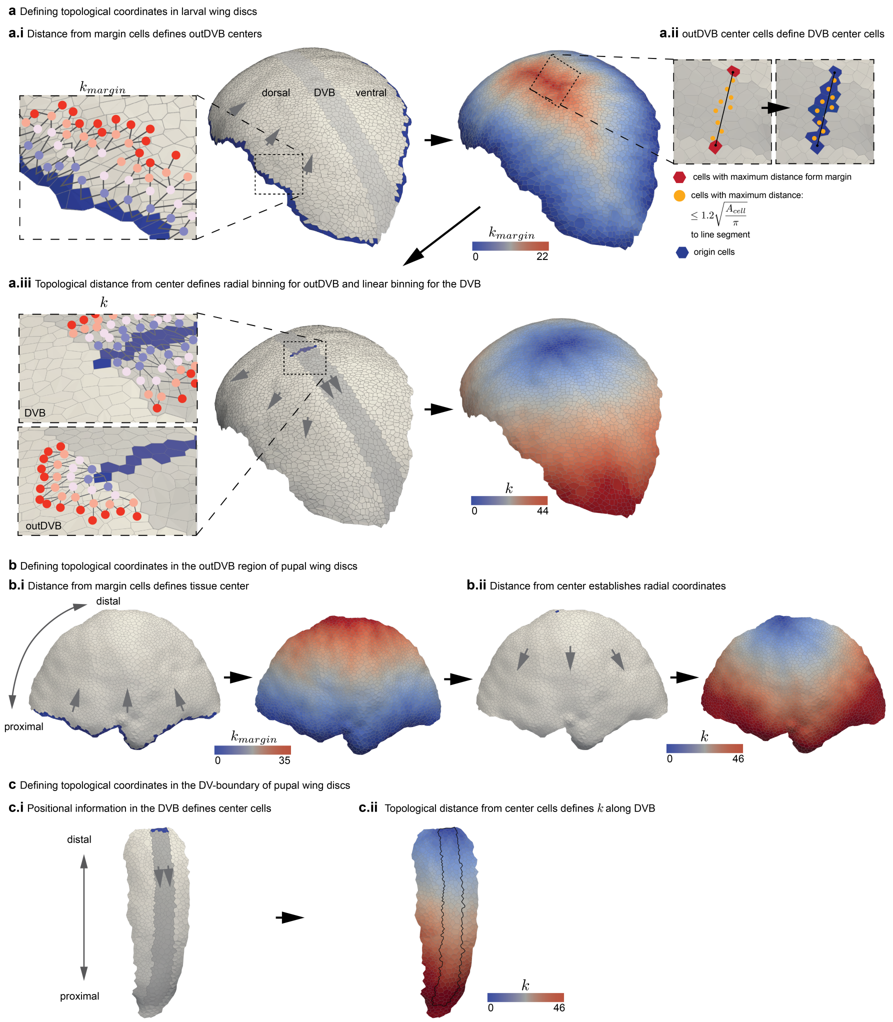


**Fig. S3 Definition of topological coordinates**: **a**, Definition of topological coordinates for larval stages. For all cells of the outDVB region, we first identify the topological distance to the outermost row of cells (blue cells, left wing pouch) in the segmentation mask that are not part of the DVB ($k_{margin}$). An example of the topology resulting from this method is shown in the inset (**a.i**). The center cells are then defined by the largest minimal $k_{margin}$ (right wing pouch and **a.ii**, red cells). This method defines a single origin cell for the topological coordinate $k$ each for the dorsal and ventral side of the outDVB (**a.iii**, lower inset). The origin for the DVB is defined as the cells lying within $\sqrt{A_{\text{cell}}/\pi} * 1.2$ distance to a straight line connecting the dorsal and ventral center cells **a.ii**. Orange circles indicate the centers of those cells that fulfill this criterion (**a.ii**, inset, left) and will serve as origin for the DVB thereafter (**a.ii**, inset, right). This method creates a line of cells as origin for the DVB, and $k$ is the minimal topological distance calculated for each cell in the DVB to any origin cell in the DVB (**a.iii**, upper inset). **b-c**, Topological coordinates for pupal stages are generated slightly differently, as pupal stages are imaged in 4 imaging angles that are 90° apart. **b**, For each cell in the outDVB region, $k_{margin}$ is found by the topological distance to the outermost row of cells in the segmentation mask that are located in the proximal region adjacent to the HP-fold (**b.i**, dark blue cells). A single origin cell is then defined by the largest minimal $k_{margin}$ (**b.ii**, dark blue cell). For the DVB, the origin is a line of cells in the DVB that is located at the most distal end of the segmentation mask (**c.i-ii**).

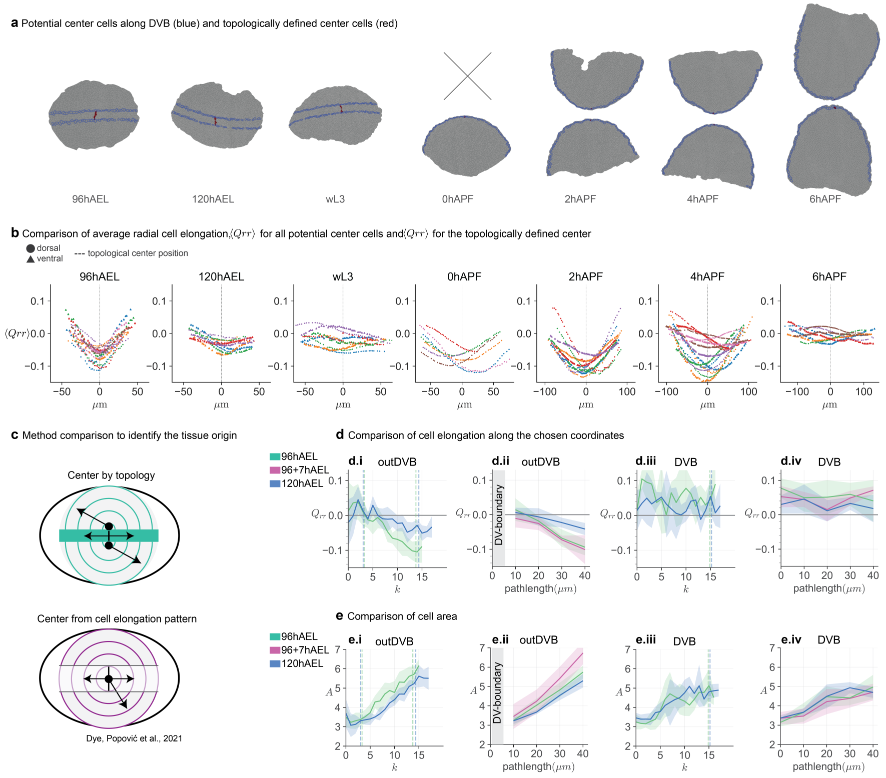


**Fig. S4 Evaluation of the topological center definition and comparison of cell elongation analysis with previous methodology**: **a**, Exemplary segmentation masks for all stages, highlighting the cells along the DVB that could qualify as a potential center cell for the outDVB region in blue and the respective topologically defined center cells in red. **b**, The tissue average $Q_{rr}$ value is calculated for each potential center cell along the DVB. Each color represents one wing disc; circles are data from dorsal outDVB regions, while triangles are from ventral. The position along the boundary is plotted on the x axis, where $0\mu m$ corresponds to the position of the topologically defined center. If the position of the minimal value for $\langle Q_{rr} \rangle$ is near 0, the topological center matches the geometrical center that would be defined from the cell elongation pattern. **c**, Schematics visualizing the sightly different spatial coordinates used to quantify patterns of cell geometry. In this work, we use topology to identify a unique center cell for each of the dorsal and ventral sides and a line of center cells for the DVB (top, green). Previous work [33] used cell elongation to identify a global tissue center, which was used to define polar coordinates in the outDVB and cartesian coordinates for the DVB (bottom, magenta). **d-e**, Comparison of the cell geometry pattern in larval wing discs using the two methods described in c: i, iii, use the new topological method to calculate the cell geometry at 96hAEL (hours After Egg Laying) and 120hAEL (with 5 wing discs each); ii, iv, directly compare both methods by plotting the new data over distance in $\mu m$ together with data published in Ref [33] from explanted wing discs from 96hAEL after a culture time of 7hrs (96+7hAEL, N=5). Solid lines indicate the average over all wing discs; the shaded region indicates the standard deviation. Note that in Ref [33], the first $10\mu m$ around the DVB were removed from the outDVB region. Here, we start the topological binning next to the DVB, which we measure to be $5\mu m$. The range of 10 - $40\mu m$ for outDVB and the range from 0 - $40\mu m$ for the DVB are used for comparison. The vertical lines in i and iii indicate upper and lower limits on $k$ covering this range. Plots show the radial component of cell elongation $Q_{rr}$ (**d**) and cell area $A$ (**e**) over $k$ and over distance to the origin in $\mu m$ in the outDVB and DVB, respectively. The data for 96+7hAEL are taken from Ref [33].

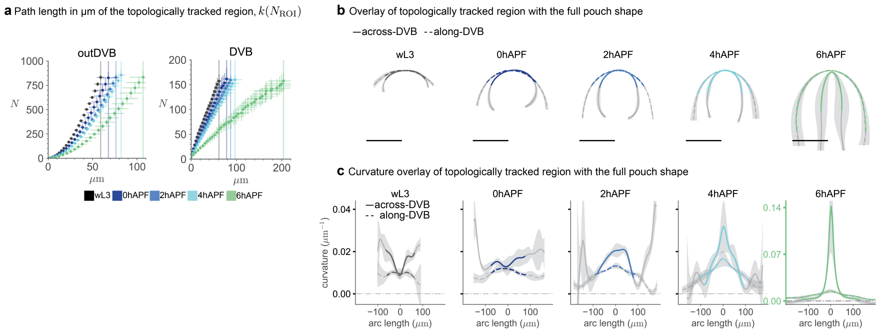


**Fig. S5 Tissue curvature and shape quantification in the topologically tracked region**: **a**, The number of cells ($N$) contained within $k$ is plotted over the path length from the origin in $\mu m$. The path length is the shortest path along cell centers from each cell to the origin. Each datapoint corresponds to the average $N$ and path length ($\mu m$), with bars indicating the standard deviation. The vertical lines indicate the average distance from origin of the maximum $k$ for each stage. **b**, Mean (solid line) and standard deviation (ribbon) of the pouch shape in the across-DVB and along-DVB directions for minimum 5 wing discs per developmental stage. The region covered by the topologically tracked region overlaid in color on the full pouch shape, with color indicating time. We use the average path length at largest $k$ in the topologically tracked region ($k(N_{\mathrm{ROI}})$) to define the size of this region. For the across-DVB cross-section, we increase this region by the average width of the DVB for each stage. **c**, Mean curvature (solid line) and standard deviation (ribbon) over arc length for the pouch ($\mu m$); the region covered in maximum $k$ is overlaid in color, with color indicating time. For the curvature and shape plots, anterior is left in the along-DVB direction, and dorsal is left for the across-DVB direction. The y-axis scale is the same through wL4 to 4hAPF. Scale bars = $100\mu m$.

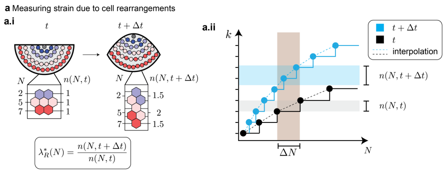


**Fig. S6 Quantification of strain due to rearrangements from static data**: **a**, Schematics explaining the quantification of strain due to cell rearrangements. **a.i** Partial wing disc pouches are shown at developmental stages $t$ and $t + \Delta t$. Cells are shown by circles, colored by the topological ring in which they lie at stage $t$. A smaller patch is shown in the inset, where $N$ denotes the cumulative number of cells from an origin until each topological ring. For example, the second topological ring at developmental stage $t$ in the inset has $N = 5$, since it has $\Delta N = 3$ cells and the ring above it has 2 cells. Then, we define $n(N, t)$ to be the measure of the number of rings needed to contain $\Delta N$ cells. $n(5, t) = 1$ (by construction); $n(5, t + \Delta t) = 2$, since at developmental stage $t + \Delta t$, the cells from this ring end up forming one complete topological ring and two half topological rings. The strain due to rearrangements at location $N$ is defined as the ratio $\lambda^*_R(N) = n(N, t + \Delta t)/n(N, t)$. **a.ii**, Plot of the topological ring number $k$ versus the cumulative cell number $N$. We focus on a single ring containing $\Delta N$ cells, visualized by the brown rectangle. $n(N, t)$ and $n(N, t + \Delta t)$ are calculated by taking the difference between the $k$ values at which the brown rectangle intersects with the interpolated $k$-curve (black dashed line for $t$ and blue dashed line for $t + \Delta t$). See also Methods 7.12.

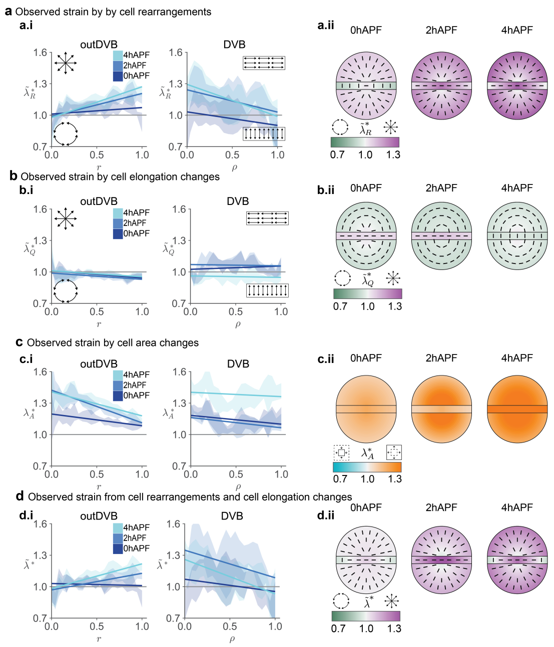


**Fig. S7 Observed strain from different cell behaviors**: **a.i-d.i**, Observed strain from cellular behaviors as a function of normalized distance from origin $r$ and $\rho$. Measured strains arise from: rearrangements ($\tilde{\lambda}^*_R$, **a.i**); cell elongation changes ($\tilde{\lambda}^*_Q$, **b.i**); cell area changes ($\lambda^*_A$, **c.i**); rearrangements ($\tilde{\lambda}^*_R$) and cell elongation changes ($\tilde{\lambda}^*_Q$) put together (**d.i**). Visual representation of the same data on the right (**a.ii - d.ii**). Half circles indicate the outDVB region; the rectangular box indicates the DVB. The color represents the magnitude of different strains. The bars visualize the direction of anisotropic strains ($\tilde{\lambda}^*_R$, $\tilde{\lambda}^*_Q$ and $\tilde{\lambda}^*$).

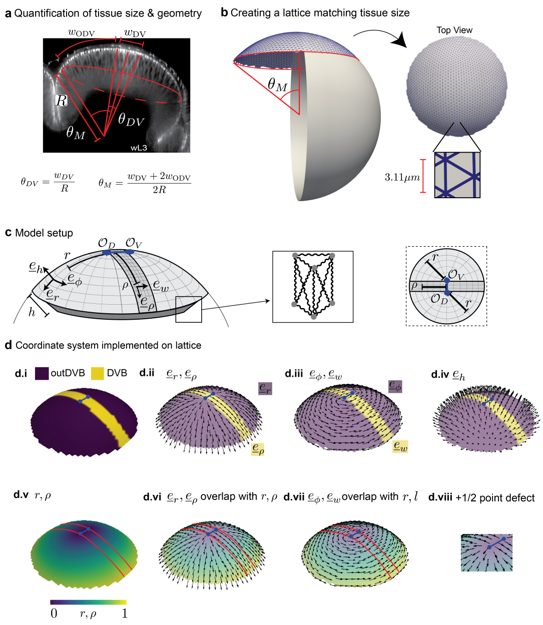


**Fig. S8 Adaption of the programmable spring model to the wing pouch geometry**: **a**, Schematic showing how the size of the wL3 wing disc pouch is calculated. $w_{DV}$ corresponds to the width of the DVB; $w_{ODV}$ corresponds to the width of the outDVB region; $R$ corresponds to the average radius of curvature at the wL3 stage. These values are used to calculate the angle that the DVB makes with the center $\theta_{DV}$ and the semi-angle of the estimated spherical cap, $\theta_M$. **b**, A spherical mesh of radius $R$ from which a spherical mesh of semi-angle $\theta_M$ is cropped. The top view of the lattice shows the spatial distribution of points on the spherical surface. The zoom-in shows the average length of the in-plane spring lattice. **c**, Imposing a coordinate system in the model similar to the coordinate system observed in the data. The model is divided into outDVB regions with a DVB region in the middle. In the outDVB regions, we have origins as $\mathcal{O}_D$ and $\mathcal{O}_V$ and coordinate bases as $(\underline{e}_r, \underline{e}_\phi, \underline{e}_h)$. In the DVB region, we have a center line $\mathcal{O}_{DV}$ and coordinate bases $(\underline{e}_\rho, \underline{e}_w, \underline{e}_h)$. We define a radial coordinate in the outDVB region as $r$ and a scalar coordinate $\rho$ in the DVB region. **d.i-iv**, A spherical cap mesh is shown with different basis vectors that change based on DVB or outDVB. **d.v**, Mesh colored by the scalar coordinates $r$ and $\rho$. **d.vi-vii**, Mesh showing how the coordinate basis vectors change with $r$ and $\rho$. **d.viii** Zoom-in of the mesh showing how our coordinate system introduces a $+1/2$ point defect at $O_D$ and $O_V$.

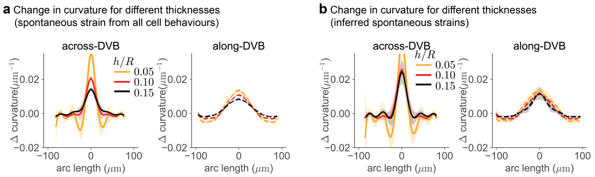


**Fig. S9 Thickness sweep to infer the bending rigidity**: **a**, Resulting change in curvature for simulations in which spontaneous strains come from the observed patterns of all cell behaviors combined ($\underline{\underline{\lambda}} = \{\lambda^*_A, \tilde{\lambda}^*_Q \tilde{\lambda}^*_R\}$) and thickness is varied. $h$ is the thickness of the model; $R$ is the radius of curvature of the top surface of the model. Simulation with $h/R = 0.10$ best matches the observed change in curvature from wL3 to 4hAPF in the data on the wing disc pouch (Fig. 4a). **b**, Resulting change in curvature for simulations with inferred spontaneous strains from active cell behaviors ($\underline{\underline{\lambda}} = \{\lambda^*_A/\lambda^{\rm res}_R, \tilde{\lambda}^*_R\}$) and varying thickness.

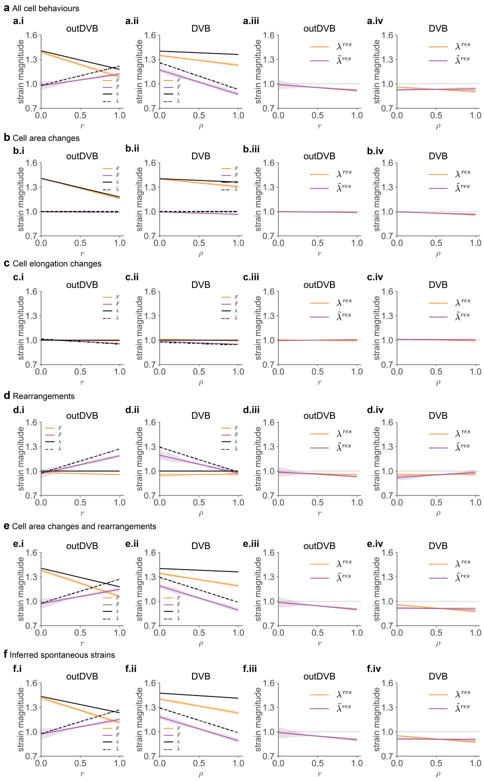


**Fig. S10 Input spontaneous strain and residual strain in *wild type* simulations for different cell behaviors**: **a - f**, Magnitudes of input spontaneous strains for *wild type* simulations are shown plotted against the radial coordinate $r$ (outDVB, **a.i - f.i**) and $\rho$ (DVB, **a.ii - f.ii**). The isotropic component of input spontaneous strain ($\lambda$) is plotted as a solid black line, and the anisotropic component of input spontaneous strain ($\tilde{\lambda}$) is plotted as a dashed black line. The isotropic component of resulting strain ($F$) in the simulation is plotted as the solid orange line, and the anisotropic component of resulting strain ($\tilde{F}$) in the simulation is plotted as the solid purple line. The magnitude of residual strain is plotted against the radial coordinate $r$ (outDVB, **a.iii - f.iii**) and $\rho$ (DVB, **a.iv - f.iv**). The isotropic component of residual strain ($\lambda^{\mathrm{res}}$) is plotted as the solid orange line, and the anisotropic component of residual strain ($\tilde{\lambda}^{\mathrm{res}}$) is plotted as the solid purple line.

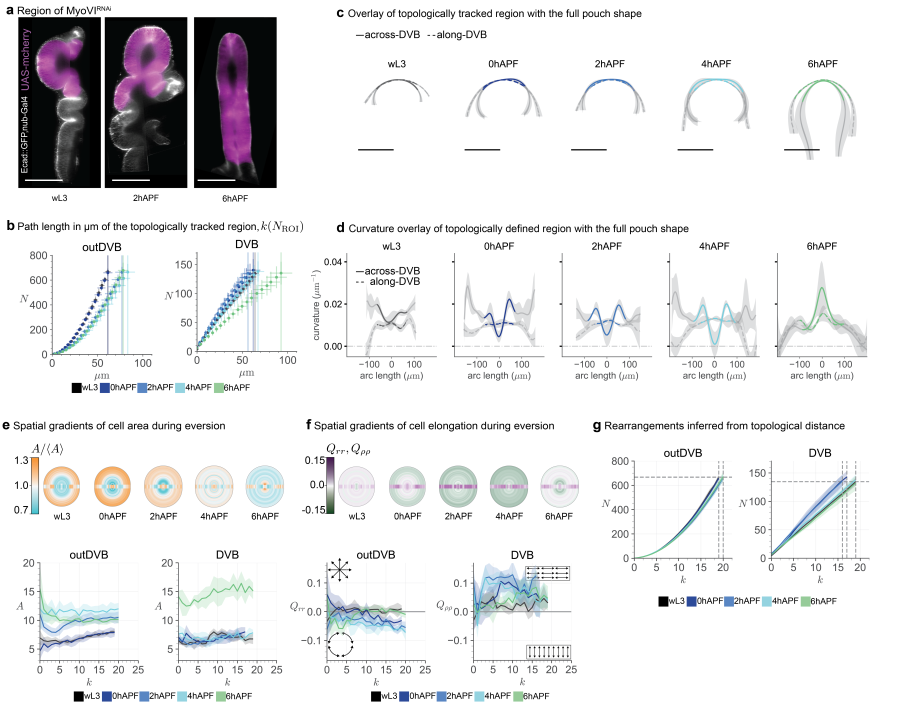


**Fig. S11 Quantification of tissue shape and cell behaviors in MyoVI$^{\mathbf{RNAi}}$**: **a**, The UAS-mCherry construct was used to visualize the expression domain of nub-Gal4, which was later used to knockdown MyoVI using RNAi. **b-g**, Analysis of the MyoVI$^{\mathrm{RNAi}}$ phenotype. **b**, The cumulative number of cells ($N$) contained up to $k$ over the path length from origin in $\mu m$. Each data point corresponds to the average $N$ and path length ($\mu m$) with its standard deviation. The vertical lines indicate the average distance from origin of the maximum $k$ for each stage. **c**, Mean $xy$ position and standard deviation of the pouch shape in the across-DVB and along-DVB direction for minimum 5 wing discs per developmental stage. The region covered by the average distance from origin for maximum $k$ is overlaid in color on the full pouch shape. For the across-DVB cross-section, we increase this region by the average width of the DVB for each stage. **d**, Mean curvature and standard deviation over arc length for the pouch ($\mu m$); the region covered in maximum $k$ is overlaid in color. **e,f**, Cell shape properties spatially averaged between wing discs and k; Dorsal and ventral are averaged together into 'outDVB'. Spatial representations show the outDVB region as half-circles; the DVB is visualized by a rectangular box. **e**, Spatial representations show the cell area gradients within each stage. x,y plots show the average cell area over the topological distance k, with the 95% confidence of the mean. **f**, Cell elongation along $Q_{rr}$ and $Q_{\rho\rho}$ for each stage; color coded in the spatial representations and plotted over k in the x,y plots with the 95% confidence of the mean. **g**, Cumulative number of cells $N$ over $k$. The horizontal line shows the maximum $N$ for the largest topological ring at wL3 stage; the vertical lines show the $k$ at which this number of cells is contained per time point. Scale bars = 100$\mu m$.

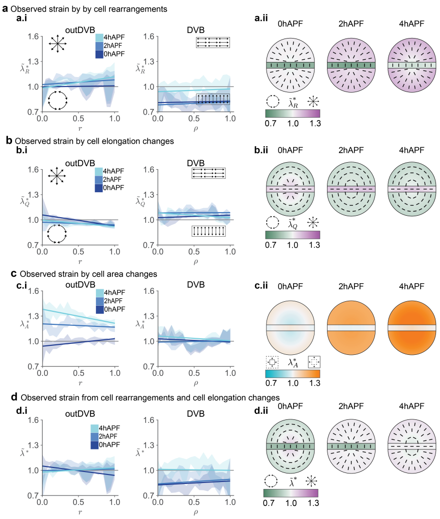


**Fig. S12 Observed strain from different cell behaviors in MyoVI$^{\mathrm{RNAi}}$: a.i-d.i**, Observed strain from cellular behaviors in the MyoVI$^{\mathrm{RNAi}}$ wing disc as a function of normalized distance from origin $r$ and $\rho$. Measured strains arise from: rearrangements ($\tilde{\lambda}^*_R$, **a.i**), cell elongation changes ($\tilde{\lambda}^*_Q$, **b.i**), cell area changes ($\lambda^*_A$, **c.i**), or rearrangements ($\tilde{\lambda}^*_R$) and cell elongation changes ($\tilde{\lambda}^*_Q$) put together (**d.i**). Visual representation of the same data on the right (**a.ii - d.ii**). Half circles indicate the outDVB region, and the rectangular box indicates the DVB. The color represents the magnitude of different $\lambda$s; the bars visualize the direction of spontaneous strain for $\tilde{\lambda}^*_R$ and $\tilde{\lambda}^*_Q$.

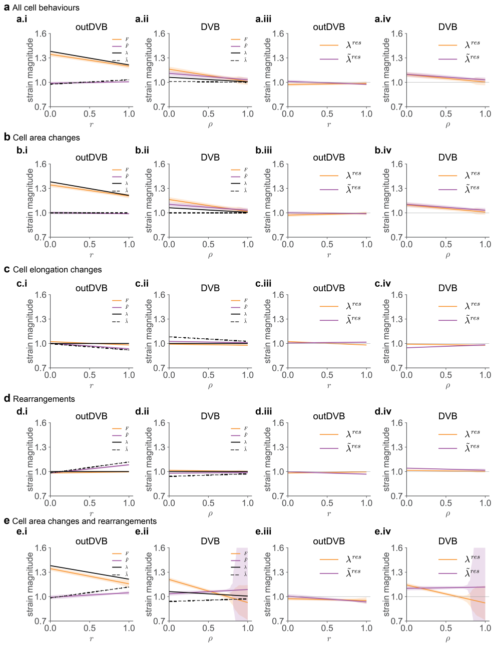

**Fig. S13 Input spontaneous strain and residual strain in MyoVI$^{\mathbf{RNAi}}$ simulations for different cell behaviors**: **a - f**, Magnitudes of input spontaneous strains for MyoVI$^{\mathrm{RNAi}}$ simulations are shown against the radial coordinate $r$ (outDVB, **a.i - e.i**) and $\rho$ (DVB, **a.ii - e.ii**). The isotropic component of input spontaneous strain ($\lambda$) is plotted as a solid black line, and the anisotropic component of input spontaneous strain ($\tilde{\lambda}$) is plotted as a dashed black line. The isotropic component of resulting strain ($F$) in the simulation is plotted as the solid orange line, and the anisotropic component of resulting strain ($\tilde{F}$) in the simulation is plotted as the solid purple line. The magnitude of residual strain is plotted against the radial coordinate $r$ (outDVB, third plot from left) and $\rho$ (DVB, **a.iv - e.iv**). The isotropic component of residual strain ($\lambda^{\mathrm{res}}$) is plotted as a solid orange line, and the anisotropic component of residual strain ($\tilde{\lambda}^{\mathrm{res}}$) is plotted as the solid purple line.

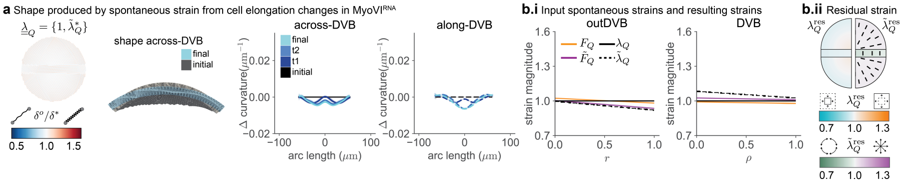


**Fig. S14 Observed in-plane in MyoVI$^{\mathbf{RNAi}}$ from cell elongation changes is inserted in the model as spontaneous strain**: **a**, The observed in-plane strain in MyoVI$^{RNAi}$ from cell elongation changes ($\tilde{\lambda}_Q^*$) is inserted in the model as spontaneous strain by a change in rest length of the springs ($\delta^o/\delta^*$). The top down view (left) shows $\delta^o/\delta^*$ from the initial to final stage; the model cross-section shows the shape in the across-DVB direction. The change in curvature of the model outcomes are plotted for all time points (right) in the across-DVB and along-DVB directions. The initial shape is a spherical cap with a radius matching the *wild type* wL3 stage. t1,t2, and final stages are the model results according to $\delta^o/\delta^*$ from the change in strains by 0,2, and 4hAPF. $\underline{\underline{\lambda}}_Q$ contains anisotropic component $\tilde{\lambda}_Q$ as observed strains from cell elongation changes $\tilde{\lambda}_Q^*$, while the isotropic component $\lambda_Q$ is set to 1. **b.i**, Input spontaneous strain for $\tilde{\lambda}_Q^*$ at the final eversion time point ($\lambda_Q$, $\tilde{\lambda}_Q$) and the resulting strain that is achieved after relaxation of the model, which can be isotropic ($F_Q$) and anisotropic ($\tilde{F}_Q$). **b.ii**, Residual strain that remains at the final time point. The color shows the magnitude of strain, and the bars the direction. The plot is split vertically to show the isotropic component ($\lambda_Q^{\text{res}}$) on the left and the anisotropic component ($\tilde{\lambda}_Q^{\text{res}}$) on the right.

**Supplemental movie 1**: Volumetric segmentation of the DP before eversion (left, wL3) and after bilayer formation (right, 2hAPF). The wing discs rotate around the across-DVB axis. The position of the pouch (blue) and the DVB (dashed line) are highlighted in two angles (top down in the first frame and anterior-lateral after 1/4 of the movie). For the wL3 wing disc ventral is up (above the DVB) and dorsal down, the first frame shows the apical side of the DP where anterior is left. For the 2hAPF wing disc, the first frame shows the apical side of the dorsal half (anterior is left) and the ventral side comes into view as the disc rotates.

**Supplemental movie 2**: Simulation with input as inferred spontaneous strain for *wild type*. Across-DVB (left) and along-DVB (right) cross-sections are shown. The initial configuration is shown in gray. The "current" configuration refers to the mesh configuration as the spontaneous strains relax (shown in blue). The initial configuration is symmetrical in the two cross-sections, while the final configuration is sharper in the across-DVB cross-section than the along-DVB cross-section.

**Supplemental movie 3**: Simulation with input as spontaneous strain due to cell area change in MyoVI$^{RNAi}$. Across-DVB (left) and along-DVB (right) cross-sections are shown. The initial configuration is shown in gray. The "current" configuration refers to the configuration that the mesh takes as the spontaneous strains relax (shown in blue). The model exhibits slight flattening in the center of the mesh in the across-DVB cross-section at the final stage. The configuration in the along-DVB cross-section does not undergo a major shape change except an overall size increase.
